# NMDA Receptor Dysregulation by Defective Depalmitoylation in the Infantile Neuronal Ceroid Lipofuscinosis Mouse Model

**DOI:** 10.1101/390732

**Authors:** Kevin P Koster, Walter Francesconi, Fulvia Berton, Sami Alahmadi, Roshan Srinivas, Akira Yoshii

## Abstract

Protein palmitoylation and depalmitoylation alter protein function. This post-translational modification is critical for synaptic transmission and plasticity. Mutation of the depalmitoylating enzyme palmitoyl-protein thioesterase 1 (PPT1) causes infantile neuronal ceroid lipofuscinosis (CLN1), a pediatric neurodegenerative disease. However, the role of protein depalmitoylation in synaptic maturation is unknown. Therefore, we studied synapse development in *Ppt1^-/-^* mouse visual cortex. We demonstrate the stagnation of the developmental N-methyl-D-aspartate receptor (NMDAR) subunit switch from GluN2B to GluN2A in *Ppt1^-/-^* mice. Correspondingly, GluN2A-mediated synaptic currents are diminished and *Ppt1^-/-^* dendritic spines maintain immature morphology *in vivo*. Further, GluN2B is hyperpalmitoylated in *Ppt1^-/-^* neurons and associated with extrasynaptic, diffuse calcium influxes and enhanced vulnerability to NMDA-induced excitotoxicity. Remarkably, *Ppt1^-/-^* neurons treated with palmitoylation inhibitors demonstrate normalized levels of palmitoylated GluN2B and Fyn kinase, reversing susceptibility to excitotoxic insult. Thus, depalmitoylation of GluN2B by PPT1 plays a critical role in postsynapse maturation and pathophysiology of neurodegenerative disease.

## Introduction

The neuronal ceroid lipofuscinoses (NCLs) are a class of individually rare, primarily autosomal recessive, neurodegenerative diseases occurring in an estimated 2 to 4 of 100,000 live births (Nita et al., 2016). Collectively, NCLs represent the most prevalent class of hereditary pediatric neurodegenerative disease (Haltia, 2006). The NCLs are characterized by progressive neurodegeneration, blindness, cognitive and motor deterioration, seizures, and premature death. The cardinal feature of all NCLs is the intracellular accumulation of proteolipid material, termed lipofuscin (Jalanko and Braulke, 2009; Nita et al., 2016). While lipofuscin accumulates in all cells of affected individuals, it deposits most robustly in neurons. This accumulation is concurrent with rapid and progressive neurodegeneration, particularly of thalamic and primary sensory cortical areas (Bible et al., 2004; Kielar et al., 2007). The NCLs are categorized into *CLN1-14* based on the age of onset and the causative gene mutated. The products of *CLN* genes are lysosomal and endosomal proteins, therefore NCLs are also classified as lysosomal storage disorders (LSDs) (Bennett and Hofmann, 1999; Jalanko and Braulke, 2009). The infantile form of disease, CLN1, presents as early as 6 months of age with progressive psychomotor deterioration, seizure, and death at approximately five years of age (Haltia, 2006; Jalanko and Braulke, 2009; Nita et al., 2016). CLN1 disease is caused by mutations in the gene *CLN1*, which encodes the enzyme palmitoyl-protein thioesterase 1 (PPT1) (Camp and Hofmann, 1993; Camp et al., 1994; Vesa et al., 1995; Jalanko and Braulke, 2009). PPT1 is a depalmitoylating enzyme responsible for the removal of palmitic acid from modified proteins (Camp and Hofmann, 1993; Lu and Hofmann, 2006).

Protein palmitoylation, the addition of a 16-carbon fatty acid (palmitic acid) to cysteine residues, is a crucial regulator of protein trafficking and function, particularly in neurons (Hayashi et al., 2005; Kang et al., 2008; Noritake et al., 2009; Fukata and Fukata, 2010; Fukata et al., 2013; Han et al., 2015). This post-translational modification is mediated by palmitoyl acyltransferases (PATs) of the zDHHC enzyme family (Fukata et al., 2006; Fukata and Fukata, 2010; Korycka et al., 2012). In contrast to other types of protein acylation, palmitoylation occurs via a reversible thioester bond (s-palmitoylation), permitting dynamic control over target protein interactions and function. Further, palmitoylated proteins require depalmitoylation prior to lysosomal degradation (Lu et al., 1996; Lu and Hofmann, 2006). Consequently, protein palmitoylation and depalmitoylation contribute significantly to mechanisms underlying synaptic plasticity and endosomal-lysosomal trafficking of proteins (Hayashi et al., 2005, 2009; Lin et al., 2009; Noritake et al., 2009; Fukata and Fukata, 2010; Thomas et al., 2013; Keith et al., 2012; Thomas et al., 2012; Fukata et al., 2013; Woolfrey et al., 2015; Han et al., 2015; Kaur et al., 2016). Indeed, PPT1 is a lysosome-targeted depalmitoylating enzyme that localizes to the axonal and synaptic compartments (Verkruyse and Hofmann, 1996; Ahtiainen et al., 2003; Kim et al., 2008). The synaptic association of PPT1 and prominence of palmitoylated synaptic proteins suggests that PPT1 influences synaptic functions through, at least, protein turnover. Many synaptic proteins undergo palmitoylation, including, but not limited to: postsynaptic density protein 95 (PSD-95), all GluA subunits of AMPARs, and the GluN2A/2B subunits of NMDARs (Kang et al., 2008). However, the role of depalmitoylation in regulating synaptic protein function remains less clear.

N-methyl-D-aspartate receptors (NMDARs) are voltage-dependent, glutamate-gated ion channels consisting of two obligatory GluN1 subunits and two GluN2 subunits that undergo a developmental change (Cull-Candy et al., 2001; van Zundert et al., 2004; Lau and Zukin, 2007; Paoletti et al., 2013). NMDARs play a crucial role in synaptic transmission, postsynaptic signal integration, synaptic plasticity (Cull-Candy et al., 2001; Van Dongen, 2009; Paoletti et al., 2013), and have been implicated in various neurodevelopmental and psychiatric disorders (Lau and Zukin, 2007; Lakhan et al., 2013; Yamamoto et al., 2015; Hu et al., 2016). NMDAR subunit composition, receptor localization, and downstream signaling mechanism undergo developmental regulation (Watanabe et al., 1992; Monyer et al., 1994; Sheng et al., 1994; Li et al., 1998; Stocca and Vicini, 1998; Tovar and Westbrook, 1999; Losi et al., 2003; van Zundert et al., 2004; Paoletti et al., 2013; Wyllie et al., 2013). In particular, GluN2B-containing NMDARs, which are abundant neonatally and allow copious calcium entry, are supplanted by GluN2A-containing NMDARs in response to experience-dependent neuronal activity (Quinlan et al., 1999a, 1999b; Philpot et al., 2001; Liu et al., 2004; Paoletti et al., 2013). This developmental switch of GluN2B-to GluN2A-containing NMDARs during brain maturation is mediated by the postsynaptic scaffolding receptors, SAP102 and PSD-95, respectively; SAP102-GluN2B-NMDAR complexes are replaced by PSD-95-GluN2A-NMDAR complexes in response to developmental, experience-dependent neuronal activity (Sans et al., 2000; van Zundert et al., 2004; Elias et al., 2008). While PSD-95, GluN2B, and GluN2A all undergo palmitoylation, how depalmitoylation regulates the turnover of these proteins, let alone during the GluN2B to GluN2A subunit switch, is unclear.

In the current study, we investigated the cellular and synaptic effects of PPT1-deficiency using the *Ppt1^-/-^* mouse model of CLN1 disease. We focused on the visual system in *Ppt1^-/-^* animals for two reasons. First, cortical blindness is a characteristic feature of CLN1 disease. Second, the rodent visual system is a well-studied model of cortical development and synaptic plasticity/maturation and it therefore serves as an optimal experimental model to examine the role of PPT1-mediated depalmitoylation during development. We found that lipofuscin accumulated very early in the *Ppt1^-/-^* visual cortex, shortly after eye-opening at postnatal day (P) 14, a timing earlier than previously documented (Gupta et al., 2001). Using biochemistry and electrophysiology, we found impeded developmental NMDAR subunit switch from GluN2B to GluN2A in *Ppt1^-/-^* mice compared to wild-type (WT). This NMDAR disruption is associated with disrupted dendritic spine morphology *in vivo*. To gain further mechanistic insight into neurodegeneration in CLN1, we used cultured cortical neurons and found that *Ppt1^-/-^* neurons recapitulate the disrupted GluN2B to GluN2A switch, leading to excessive extrasynaptic calcium transients and enhanced vulnerability to NMDA-mediated excitotoxicity. We directly examined protein palmitoylation state and found hyperpalmitoylation of GluN2B as well as Fyn kinase, which facilitates GluN2B surface retention, in *Ppt1^-/-^* neurons. Finally, we demonstrate that chronic treatment of *Ppt1^-/-^* neurons with palmitoylation inhibitors normalized GluN2B and Fyn kinase hyperpalmitoylation and rescued the enhanced susceptibility to excitotoxicity. Our results indicate that PPT1 plays a critical role in the developmental GluN2B to GluN2A subunit switch and synaptic maturation. Further, our results indicate that these dysregulated mechanisms contribute to CLN1 pathophysiology and may be shared features of common adult-onset neurodegenerative diseases.

## Results

To understand synaptic dysregulation in CLN1 disease, we utilized the visual cortex of *Ppt1^-/-^* animals as a model system. The rodent visual cortex undergoes timed, experience-dependent plasticity, which has been well-characterized at the systemic, cellular, and molecular levels (Bear et al., 1990; Gordon and Stryker, 1996; Hensch et al., 1998; Quinlan et al., 1999a, 1999b; Fagiolini and Hensch, 2000; Mataga et al., 2001, 2004; Philpot et al., 2001; Desai et al., 2002; Yoshii et al., 2003; Hensch, 2005; Cooke and Bear, 2010). We examined WT and *Ppt1^-/-^* littermates at the following ages: P11, P14, P28, P33, P42, P60, P78, P120, which correspond to particular developmental events in visual cortex. In mice, P11 and P14 are prior to and just after eye opening (EO) respectively. Further, the critical period in the visual cortex peaks at P28 and closes from P33 to P42. Postnatal day 60, P78, and P120 were selected as adult time points. We determined whether experience-dependent synaptic maturation is altered during the progression of CLN1 pathology.

### Lipofuscin deposits immediately following eye opening in visual cortex of *Ppt1^-/-^* mice

Although it remains controversial whether lipofuscin is toxic to neurons or an adaptive, neuroprotective mechanism, its accumulation correlates with disease progression. Therefore, we examined lipofuscin deposition in the visual cortex as a marker of pathology onset and progression. Lipofuscin aggregates are readily visible as autofluorescent lipopigments (ALs) without staining under a confocal microscope. To examine the temporal and spatial accumulation of ALs in *Ppt1^-/-^* mice, we performed quantitative histology on the visual cortex (area V1) of WT and *Ppt1^-/-^* mice during early development. Visual cortical sections were imaged at the above-mentioned developmental time points and ALs were quantified in a laminar-specific manner. We found that ALs are detectable first at P14 in *Ppt1^-/-^* visual cortex, much earlier than previously reported (**Figure 1A, B, C**).

**Figure 1.**
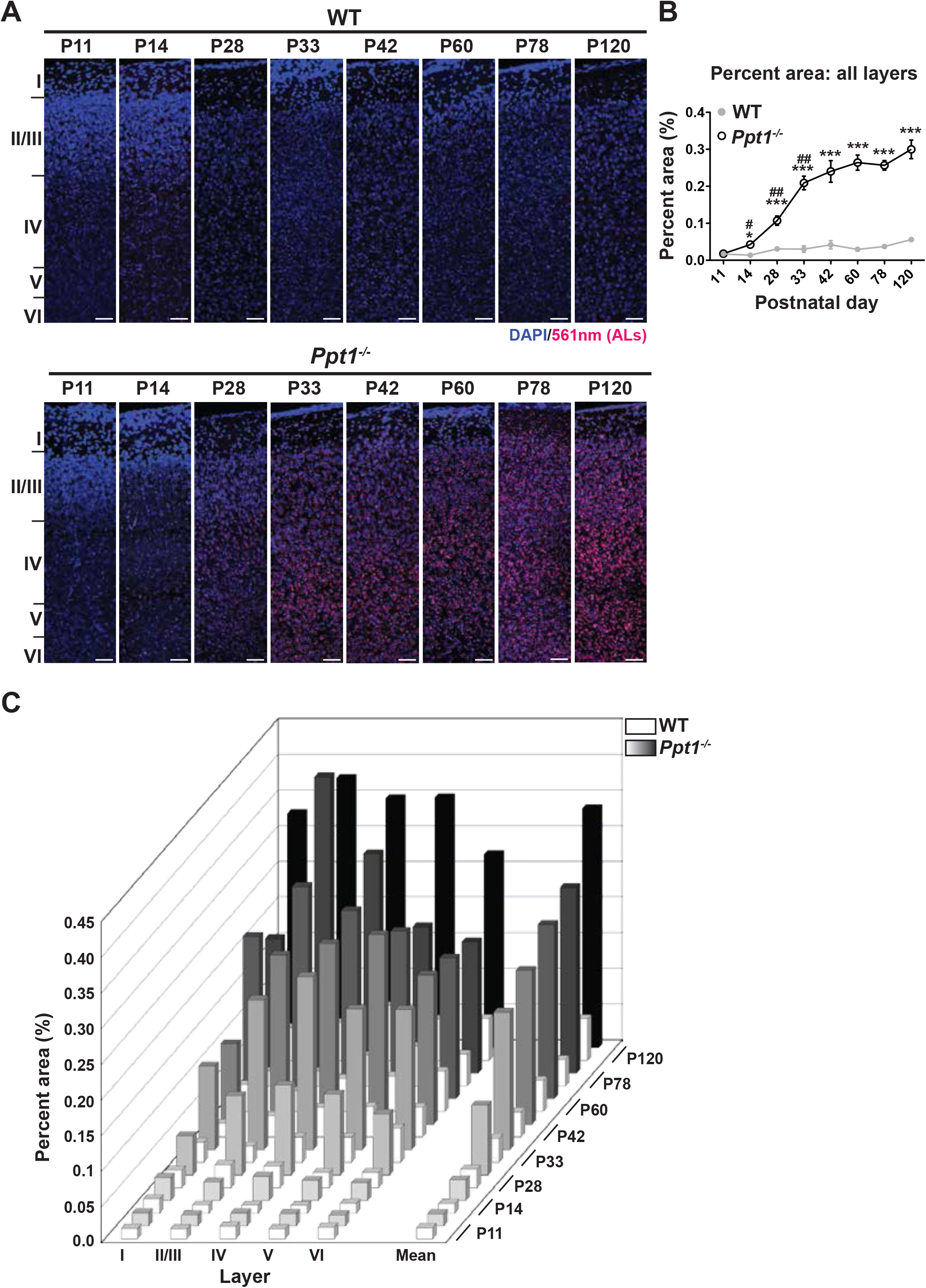
ALs deposit immediately following eye opening in visual cortex of *Ppt1^-/-^* mice. **(A)** Representative composite confocal images through area V1 of visual cortex in WT (top) and *Ppt1^-/-^* mice (bottom) during development and into adulthood. DAPI nuclear stain (blue, 405nm excitation) and AL signals (red, 561nm excitation) are visualized. Cortical layers are marked (left). Scale=50μm. **(B)** Quantification of the mean percent area occupied by ALs through all cortical layers (see methods). *Ppt1^-/-^* and WT were compared (n=4-6 animals/group) at each age using t-test and the significance was indicated as follows: *p<0.05, **p<0.01, and ***p<0.001. Significant AL increase in *Ppt1^-/-^* mouse brain between two consecutive ages (e.g. P28 vs. P33) are shown as # p<0.05 and ## p<0.01. Error bars represent s.e.m. **(C)** Cortical layer-specific quantification of area occupied by ALs separated by each cortical layer (x-axis) and age (z-axis). Averaged values, s.e.m., and n for each condition are represented in **Supplementary table 1**.

Further, AL accumulation accelerated rapidly through the critical period (Berardi et al., 2000; Hensch, 2005; Maffei and Turrigiano, 2008) and plateaued by adulthood (P60). This result indicates that neuronal AL load is saturable, and that this saturation occurs early on in disease, as *Ppt1^-/-^* animals do not perish until around 10 months old (Gupta et al., 2001). Interestingly, we noticed a trend that AL accumulation became more robust in the deep cortical layers beginning at P28, particularly layer IV, the termination site of thalamocortical neurons projecting from the dorsal lateral geniculate nucleus (dLGN) that relay retinal inputs (**Figure 1C**). WT mice accumulate a miniscule amount of ALs, even at the oldest time point examined (**Figure 1A-C**).

Whether lipofuscin accumulation is directly neurotoxic or not, profiling the temporospatial and sub-regional pattern of AL deposition will be valuable for assessing therapeutic interventions in future studies. Further, the pattern of deposition revealed herein suggests a correlation between systemic neuronal activation and lipofuscin accumulation. In particular, our result that AL deposition starts immediately following EO, the onset of patterned visual activity, and is detectable prominently in layer IV (**Figure 1A, C, Supplementary Table 1**) suggests that neuronal activity or experience-dependent plasticity are linked to lipofuscin deposition.

### NMDAR subunit composition is biased toward immaturity in *Ppt1^-/-^* visual cortex

To examine the role of PPT1 in excitatory synapse function, we focused on the NMDAR subunits, GluN2B and GluN2A, which are both palmitoylated (Hayashi et al., 2009). Developmental GluN2B to GluN2A subunit change (Paoletti et al., 2013) is critical for NMDAR function and maturation, which facilitates refinement of neural circuits and a higher tolerance to glutamate-mediated excitotoxicity (Hardingham and Bading, 2002, 2010; Hardingham et al., 2002). Furthermore, previous work shows evidence for NMDA-induced excitotoxicity in the pathogenesis of CLN1 (Finn et al., 2012). We biochemically analyzed WT and *Ppt1^-/-^* visual cortices from P11 to P60, and measured levels of GluN2B and GluN2A subunits in whole lysates and synaptosomes of WT and *Ppt1^-/-^* visual cortices. Whereas GluN2B levels were comparable between WT and *Ppt1^-/-^* at all ages, GluN2A levels in synaptosomes were significantly lower in *Ppt1^-/-^* than WT (**Figure 2A**). This decrease was present at time points during, and just following, the critical period in visual cortical development (P33, P42, and P60). When analyzed as a ratio of GluN2A/GluN2B, a robust and persistent decrease is observed in *Ppt1^-/-^* visual cortex (**Figure 2B**). GluN1 levels were unchanged between WT and *Ppt1^-/-^* in synaptosomes (**Figure 2C**), indicating the selective obstruction of GluN2A incorporation into NMDARs.

**Figure 2.**
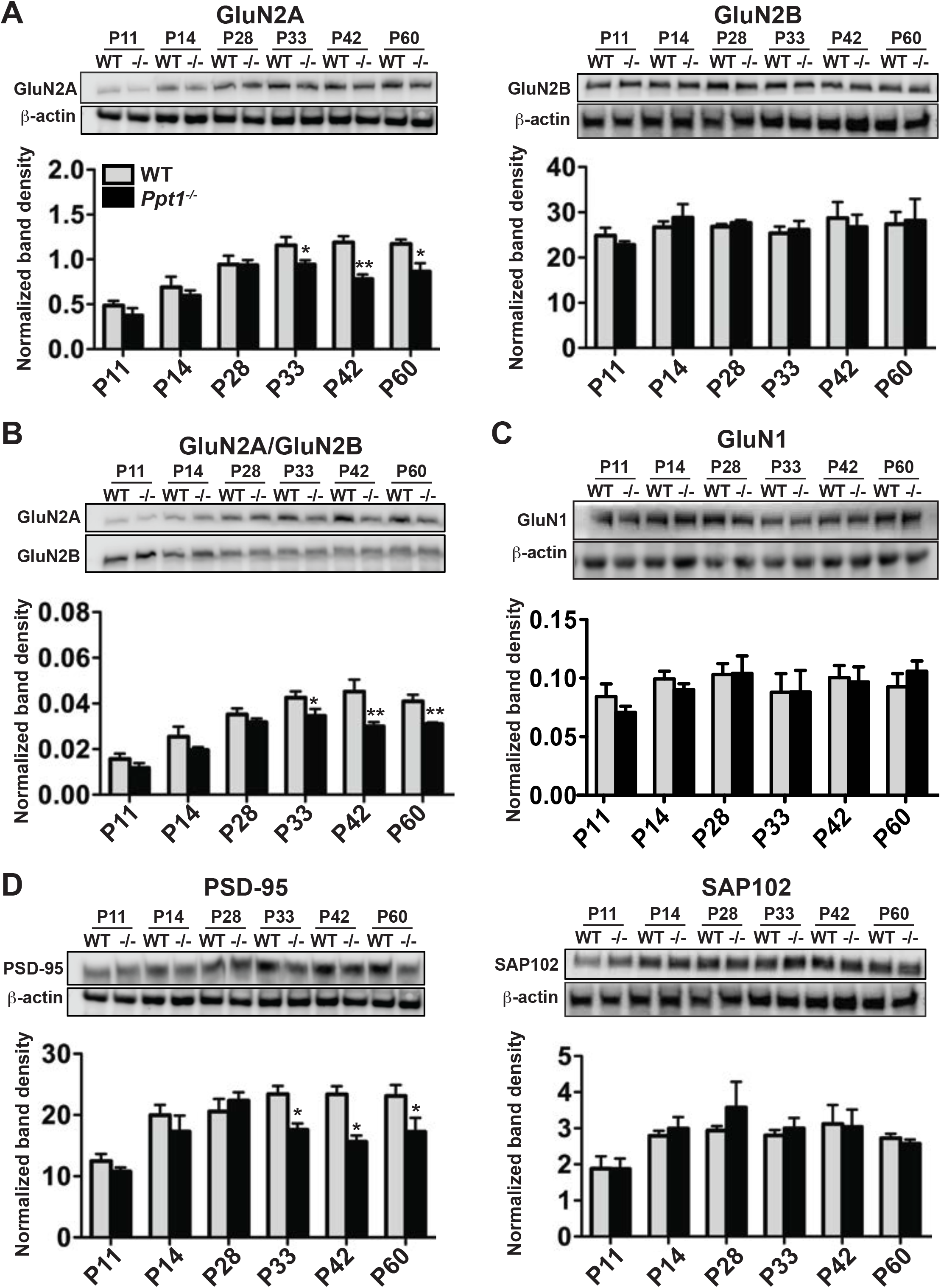
NMDAR subunit composition is biased toward immaturity in *Ppt1^-/^* visual cortex. **(A)** Representative immunoblots of GluN2 subunits, GluN2A and GluN2B across age and genotype as indicated (top) and quantification of band density (bottom) normalized to β-actin loading control within lane. **(B)** Representative immunoblots of GluN2A and GluN2B (top) and quantification of the ratio of GluN2A/GluN2B band density within animal (bottom). **(C)** Representative immunoblots of GluN1 from synaptosomes across age and genotype as indicated (top) and quantification of band density (bottom) normalized to β-actin loading control within lane. **(D)** Representative immunoblots of scaffolding molecules PSD-95 and SAP102 across age and genotype as indicated (top) and quantification of band density (bottom) normalized to β-actin loading control within lane. For all experiments in **Figure 2**, *Ppt1^-/-^* and WT were compared (n=4 independent experiments/animals with 2 repetitions/group) at each age using t-test and the significance was indicated as follows: *p<0.05, and **p<0.01. Error bars represent s.e.m.

The developmental shift from GluN2B-containing NMDARs to synaptic GluN2A-containing NMDARs is mediated by the postsynaptic scaffolding proteins, SAP102 and PSD-95 (Townsend et al., 2003; Yoshii et al., 2003; Elias et al., 2008). SAP102 preferentially interacts with GluN2B-containing NMDARs, which are enriched neonatally (Chen et al., 2000, 2011; Sans et al., 2000; Liu et al., 2004; van Zundert et al., 2004; Zheng et al., 2010). In contrast, PSD-95 has greater affinity to GluN2A-containing NMDARs, particularly in the mature brain (Sans et al., 2000; van Zundert et al., 2004; Van Dongen, 2009; Yan et al., 2014). Thus, we examined the expression of these scaffolding proteins in WT and *Ppt1^-/-^* visual cortex. Similar to the results obtained for GluN2B and GluN2A, while SAP102 levels remained unchanged, we observed a decrease in PSD-95 levels at P33-P60, the same developmental time points where GluN2A expression was reduced (**Figure 2D**). Together, these results suggest reduced incorporation and scaffolding of GluN2A-containing NMDARs in *Ppt1^-/-^* synapses, indicating immature or dysfunctional synaptic composition.

To examine whether the reduction in GluN2A is due to selective exclusion from the postsynaptic site or alterations in the total protein amount, we also measured NMDAR subunit levels in whole lysates. These findings closely match our findings in synaptosomes. Namely, GluN2A levels showed reductions in *Ppt1^-/-^* lysates beginning at the same time point (P33), while GluN2B levels were stable (**Supplementary Figure 1A**). The GluN2A/2B ratios in *Ppt1^-/-^* whole lysates were also lower than those in WT lysates and the reduction was comparable to that observed in synaptosomes. (**Supplementary Figure 1B**). GluN1 levels, however, were unaltered between genotypes (**Supplementary Figure 1C**). Collectively, these results indicate a selective decrease in the total amount of mature synaptic components in *Ppt1^-/-^* brains and suggest that synaptosomal reductions in GluN2A and PSD-95 may result from altered transcription or translation.

### NMDAR-mediated EPSCs are altered in *Ppt1^-/-^* visual cortex

Next, we sought to correlate our biochemical findings electrophysiological changes in NMDAR functionality (**Figure 2**). While human CLN1 patients present with retinal degeneration and the *Ppt1^-/-^* mouse model of CLN1 phenocopies the human disease, the electroretinogram (ERG) is effectively unaltered at 4 moths in the mouse model (Lei et al., 2006), allowing for detailed study of the electrophysiological changes in the visual cortex associated with early disease states. We recorded evoked, NMDAR-mediated excitatory postsynaptic currents (EPSCs) in layer II/III cortical neurons in visual cortical slices of WT and *Ppt1^-/-^* mice at P42. The NMDA-EPSCs were pharmacologically isolated (see Methods section) and were recorded in whole cell patch mode clamped at +50mV (**Figure 3A**). As GluN2A- and GluN2B-containing NMDARs exhibit differential receptor kinetics, with GluN2A displaying fast (~50ms) and GluN2B displaying slow decay kinetics (~300ms), their relative contribution is reliably interpolated by fitting the EPSC decay phase with a double exponential function (Stocca and Vicini, 1998; Vicini et al., 1998). From the fitting, we measured the following parameters: the amplitude (A) of the fast (Af) and slow (As) components; the ratio Af/ Af+As; the decay time constants (τ) of the fast (τf) and the slow components (τs), and the weighted decay (τw). *Ppt1^-/-^* mice showed significant decreases in the Af and ratio of Af/Af+As as compared to WT, while As showed no significant change (**Figure 3B**). Further, the τf significantly decreased in *Ppt1^-/-^* mice vs. WT, while τw and τs were comparable (**Figure 3C**). As the fast components of both the amplitude and decay constant are reduced in *Ppt1^-/-^* neurons, these data indicate a functional decrease in the contribution of GluN2A in NMDAR-mediated EPSCs in layer II/III visual cortical neurons and corroborate our biochemical findings.

**Figure 3.**
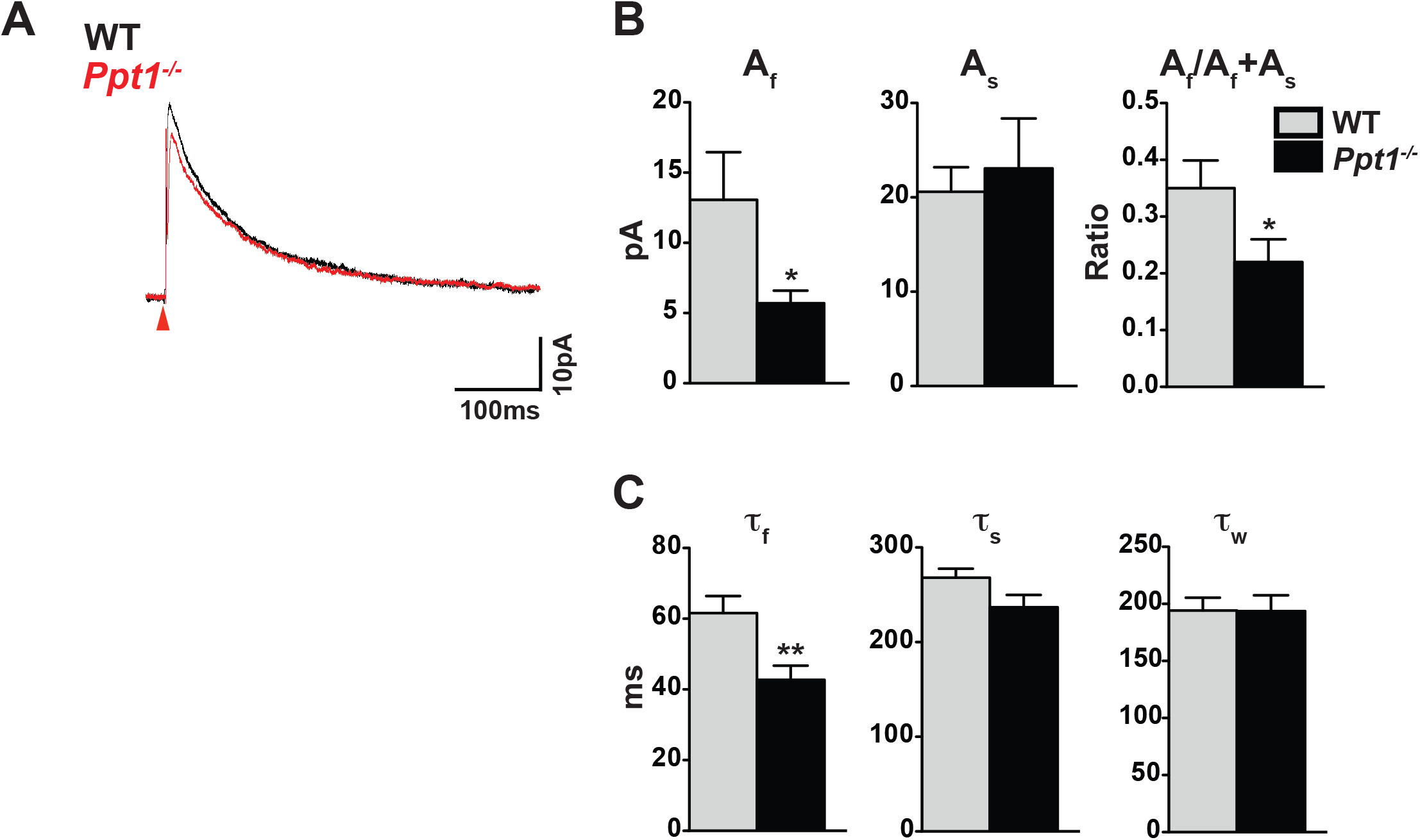
NMDAR-mediated EPSCs are altered in *Ppt1^-/-^* visual cortex. **(A)** Representative traces of NMDA-EPSCs recorded from pyramidal neurons in layer II/III of the visual cortex (V1) of WT and *Ppt1^-/-^* mice. Red arrows indicate onset of evoked stimulus. Neurons were voltage clamped at +50mV and NMDA-EPSCs evoked in layer IV. **(B)** Quantification of amplitude of the fast component, Af, slow component, As, and A_f_/A_f_+A_s_ derived from fitting the decay phase of the evoked NMDA-EPSCs with the double exponential function: Y(t) = A_f*_e ^-t/t^_fast_ + A_s*_e^-t/t^_slow_. **(C)** Quantification of the decay constant of the fast component, τf, slow component, τs, and weighted decay, τw, derived from fitting the decay phase of the evoked NMDA-EPSCs with the double exponential function: Y(t) = A_f*_e ^-t/t^_fast_ + A_s*_e^-t/t^_slow_. For experiments in **Figure 3**, *Ppt1^-/-^* and WT were compared (n=10 cells, 4 mice (WT); n=12 cells, 5 mice (*Ppt1^-/-^*)) using t-test and the significance was indicated as follows: *p<0.05, and **p<0.01. Error bars represent s.e.m.A

### Dendritic spine morphology is immature in *Ppt1^-/-^* visual cortex

The morphology of dendritic spines is dynamic, and synaptic activities directly alter spine morphology during synaptic plasticity (Engert and Bonhoeffer, 1999; Parnass et al., 2000; Matsuzaki et al., 2001; Yuste and Bonhoeffer, 2001). During visual cortical development and concomitant with the GluN2B to GluN2A switch, excitatory synaptic architecture and dendritic spine morphology undergo robust structural plasticity. In particular, dendritic spines contribute to experience-dependent synaptic plasticity via the generation, maturation, and longterm stabilization of spines, ultimately giving rise to the established synaptic circuit. Typically, by P33, dendritic spines demonstrate reduced turnover, reduced motility, and mushroom-type spine morphology, indicating synaptic maturity. Importantly, dendritic spines are morphologically disrupted in many neurodevelopmental disorders, typically skewing towards an immature phenotype (Purpura, 1979; Irwin et al., 2001; Penzes et al., 2011).

We hypothesized that dendritic spine morphology is immature or disrupted in *Ppt1^-/-^* neurons, particularly given that GluN2A subunit incorporation is disrupted *in vivo*. Thus, we used *in utero* electroporation to sparsely label layer II/III cortical neurons in visual cortex using a GFP construct (Matsuda and Cepko, 2004). GFP-expressing cells from WT and *Ppt1^-/-^* animals were imaged for detailed analysis of dendritic spine morphology (spine length, spine volume, and spine head volume) at P33, a time point when dendritic spine morphology is typically considered mature and GluN2A is reduced at *Ppt1^-/-^* synapses.

Electroporated, GFP-expressing cells (procedure schematized in **Figure 4A**) from WT and *Ppt1^-/-^* visual cortex (**Figure 4B**) were analyzed in Imaris (Bitplane) for dendritic spine characteristics. While WT neurons exhibited mushroom-type spine morphology with high-volume spine heads (**Figure 4C**, arrows), *Ppt1^-/-^* neurons showed longer, filipodial protrusions or stubby spines (**Figure 4C**, arrowheads). Quantification of spine length and spine volume demonstrated that *Ppt1^-/-^* spines were longer and less voluminous compared to WT (**Figure 4D-E**). Further, the volume of dendritic spine heads was reduced in *Ppt1^-/-^* neurons (**Figure 4E**, inset). These data indicate that dendritic spine morphology is disrupted in the developing CLN1 visual cortex, correspond with the finding that NMDAR composition is immature at P33, and suggest a reduced ability to compartmentalize calcium and other localized biochemical signals in CLN1.

**Figure 4.**
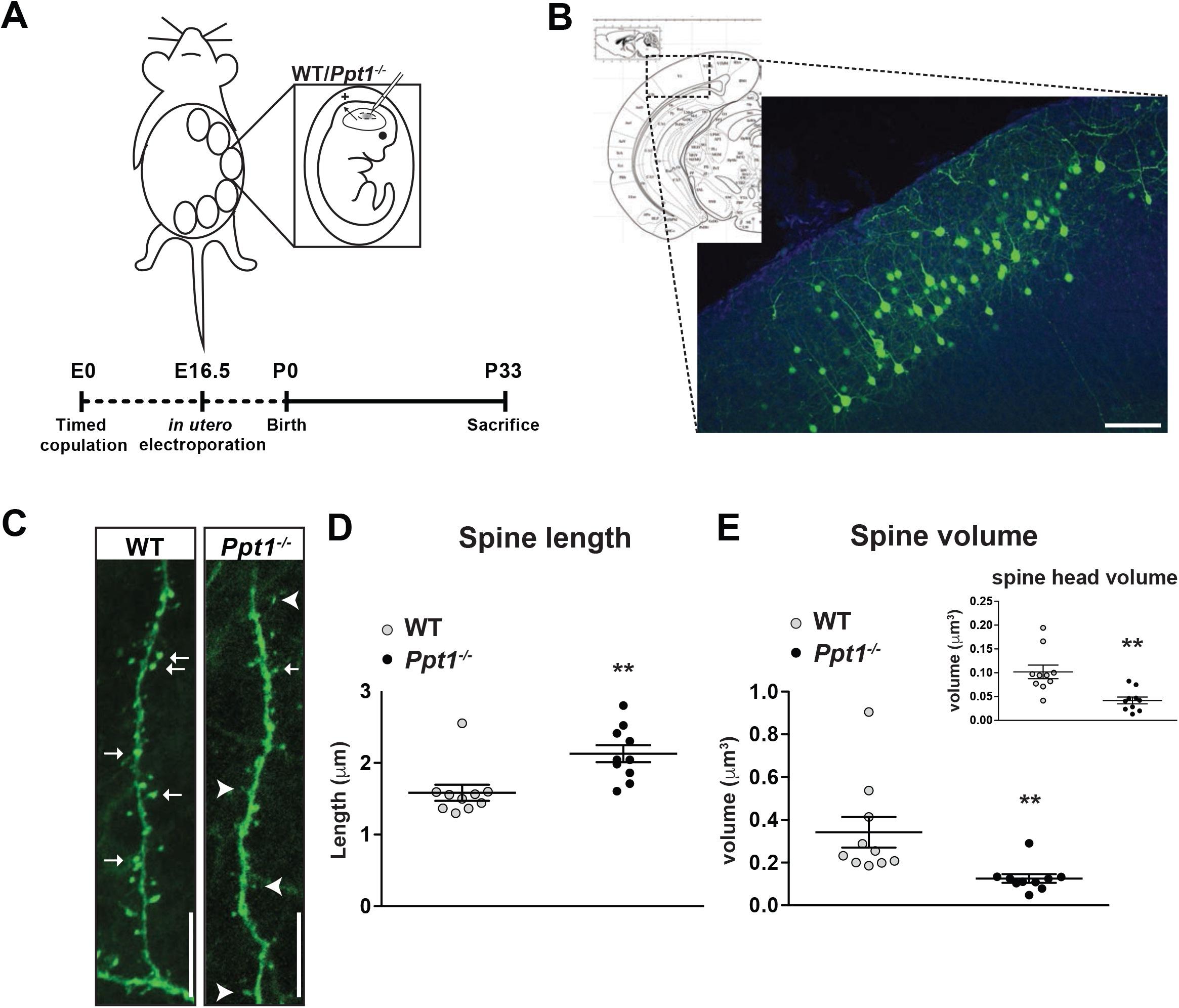
Dendritic spine morphology is immature in *Ppt1^-/-^* layer II/III visual cortical neurons. **(A)** Schematic of *in utero* electroporation procedure and timeline (bottom) **(B)** *Left*, coronal diagram from Paxino’s mouse brain atlas demonstrating areas of visual cortex and *right*, representative low-magnification (10x) confocal image of a successfully transfected group of layer II/III neurons in visual cortex. Scale bar=100μm. **(C)** Representative confocal images of GFP-transfected dendritic segments from WT and *Ppt1^-/-^* neurons at P33. Arrows mark mature, mushroom-type spines; arrowheads mark thin, filipodial spines or stubby, headless spines. Scale bar=10μm. **(D)** Semi-automated quantification of dendritic spine length in WT and *Ppt1^-/-^* visual cortical neurons at P33. **(E)** Semi-automated quantification of dendritic spine volume and spine head volume (inset) in WT and *Ppt1^-/-^* visual cortical neurons at P33. For experiments in **Figure 4**, WT and *Ppt1^-/-^* were compared (n=3-4 cells/animal, 3 animals/group) using t-test and the significance was indicated as follows: **p<0.01, *Ppt1^-/-^* vs. WT. Error bars represent s.e.m.

### NMDAR subunit composition and dendritic spine morphology are also immature in *Ppt1^-/-^* primary cortical neurons

The GluN2B to GluN2A switch and maturation of dendritic spine characteristics in WT primary neurons has been previously demonstrated (Williams et al., 1993; Zhong et al., 1994; Papa et al., 1995). We established that the developmental switch from GluN2B-to GluN2A-containing NMDARs and dendritic spine morphology are impaired in the *Ppt1^-/-^* mouse brain. To understand these mechanisms more comprehensively and examine protein palmitoylation more directly, we used dissociated neuronal cultures. First, we analyzed these developmental events in WT and *Ppt1^-/-^* primary cortical neurons to determine whether the biochemical and structural features of disease are recapitulated *in vitro*.

Cortical neurons cultured for 7, 10, or 18 days *in vitro* (DIV 7, 10, or 18) were harvested and lysates subjected to immunoblot analysis for markers of immature (GluN2B) or mature (GluN2A, PSD-95) excitatory synapses. Expression of GluN2B clearly preceded that of mature synaptic markers, peaking in both WT and *Ppt1^-/-^* neurons at DIV10 and decreasing thereafter (**Figure 5A**). In contrast, levels of both GluN2A and PSD-95 remained low until DIV18, at which point expression was robust (**Figure 5B, C**). Importantly, GluN2A, PSD-95, and GluN2A/GluN2B ratio levels were reduced in *Ppt1^-/-^* neurons compared to WT at DIV18, indicating that the biochemical phenotype is recapitulated to an extent *in vitro* (**Figure 5B-D**).

**Figure 5.**
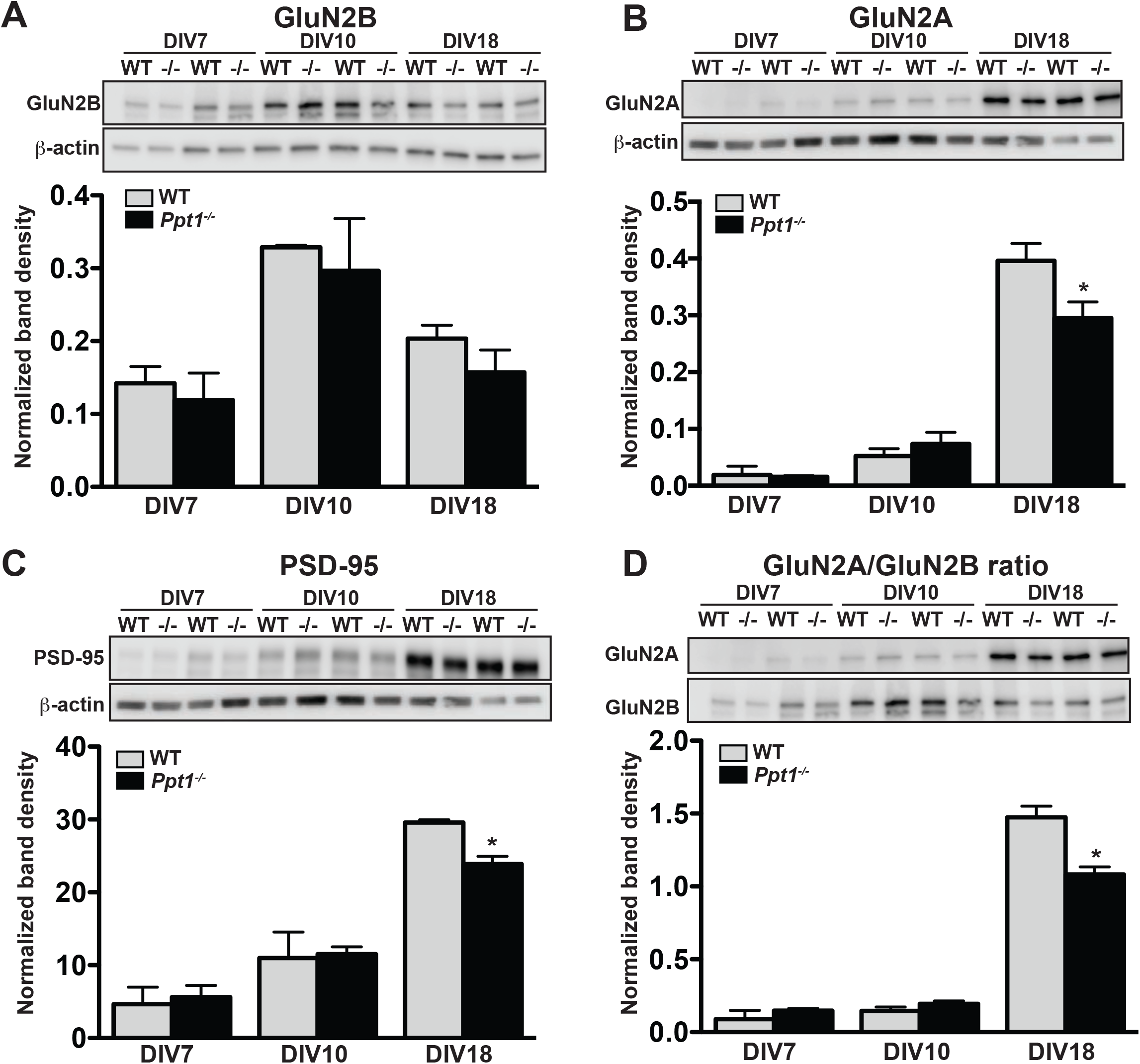
GluN2B to GluN2A NMDAR switch and Ppt1^-/-^-induced synaptic deficits are recapitulated in primary cortical neurons. **(A)** Representative immunoblot (top) and quantification of GluN2B levels in WT and *Ppt1^-/-^* neurons at DIV7, 10, and 18. **(B)** Representative immunoblot (top) and quantification of GluN2A levels (bottom) in WT and *Ppt1^-/-^* neurons at DIV7, 10, and 18. **(C)** Representative immunoblot (top) and quantification of PSD-95 levels (bottom) in WT and *Ppt1^-/-^* neurons at DIV7, 10, and 18. **(D)** Representative immunoblot (top) and quantification of the GluN2A/2B ratio (bottom) in WT and *Ppt1^-/-^* neurons at DIV7, 10, and 18. For all experiments in **Figure 5**, *Ppt1^-/-^* and WT were compared (n=2 independent experiments with 2 repetitions/group) at each time point using t-test and the significance was indicated as follows: *p<0.05 *Ppt1^-/-^* vs. WT. Error bars represent s.e.m.

To analyze dendritic spine morphology, primary cortical neurons from fetal WT and *Ppt1^-/-^* mice were transfected with a GFP construct (Matsuda and Cepko, 2004) and were cultured until DIV 15 or 20 when live cell imaging was performed (**Figure 6A**). We measured dendritic spine length and volume in transfected cells using the Imaris software (Bitplane). At both DIV 15 and 20, we observed a significant shift in the dendritic spine length and volume (**Figure 6B** and C). *Ppt1^-/-^* neurons demonstrated a significantly higher percentage of long, thin protrusions (filipodia-type; **Figure 6B**), and a significant reduction in the percentage of mature, mushroom-type dendritic spines with volumes greater than 0.2μm^3^ (**Figure 6C**). When averaged, the data demonstrate a robust increase in mean spine length and a decrease in spine volume in *Ppt1^-/-^* neurons compared to WT controls at both time points (**Figure 6B** and C, right). Together, these data demonstrate that *Ppt1^-/-^* neurons in culture give rise to morphologically immature dendritic spines and corroborate our *in vivo* findings.

**Figure 6.**
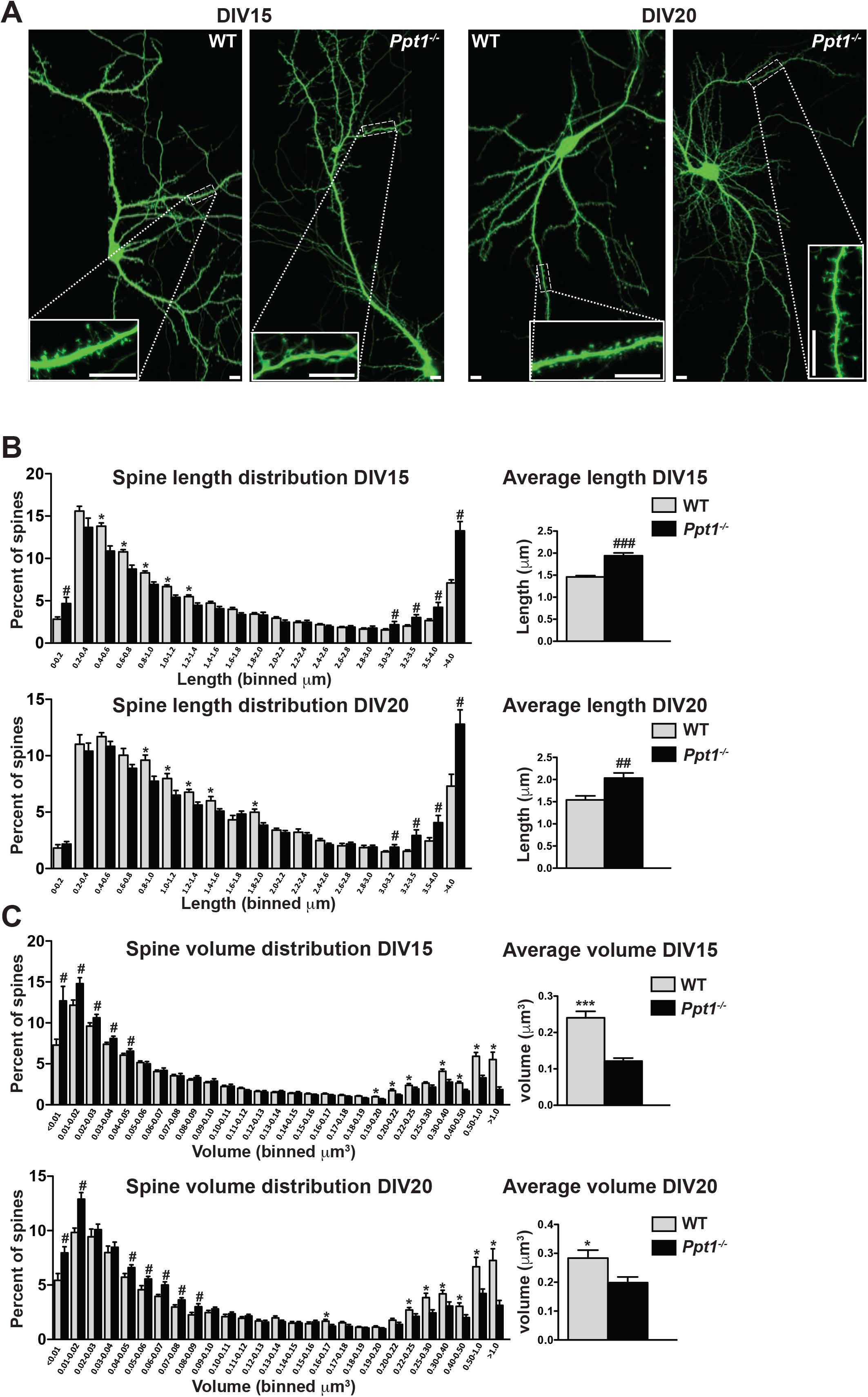
Dendritic spine morphology is immature in *Ppt1^-/-^* neurons *in vitro*. **(A)** Representative composite confocal images of live DIV15 (left) and DIV20 (right) GFP-transfected, cultured WT and *Ppt1^-/-^* neurons. Insets represent dendrite segments within dotted line. Scale bar=10μm. **(B)** *Left*, quantification of dendritic spine length in WT and *Ppt1^-/-^* neurons at DIV15 and 20. Spine length is binned into 19 discrete groups from 0 - >4μm. *Right*, mean length of all spines in cultured WT and *Ppt1^-/-^* neurons at DIV15 and 20. **(C)** *Left*, semi-automated quantification of dendritic spine volume in WT and *Ppt1^-/-^* cultured neurons at DIV15 and 20. Spine volume is binned into 27 discrete groups form 0 - >1μm^3^. *Right*, mean volume of all spines in cultured WT and *Ppt1^-/-^* neurons at DIV15 and 20. For experiments in **Figure 6**, *Ppt1^-/-^* and WT were compared (For DIV15: n=4-5 neurons/group, 3-independent experiments, WT=21,514 spines; *Ppt1^-/-^*=18,013 spines. For DIV20: n=3 neurons/group, 2-independent experiments, WT=11,335 spines; *Ppt1^-/-^*=9,958 spines) using t-test and the significance was indicated as follows: *Left*, *p<0.05, within bin where *Ppt1^-/-^* is decreased compared to WT; #p<0.05, within bin where *Ppt1^-/-^* is increased compared to WT. *Right*, ##p<.0.01 and ###p<0.001 where *Ppt1^-/-^* is increased compared to WT; *p<0.05, ***p<0.001 where *Ppt1^-/-^* is decreased compared to WT. Error bars represent s.e.m.

### Calcium imaging reveals extrasynaptic calcium dynamics in *Ppt1^-/-^* neurons

Intracellular calcium dynamics, compartmentalization, and signaling play a critical role in synaptic transmission and plasticity. These properties are altered by glutamate receptor composition and location (Lau and Zukin, 2007; Hardingham and Bading, 2010; Paoletti et al., 2013). GluN2B-containing NMDARs maintain a prolonged open conformation compared to GluN2A-containing receptors, allowing increased calcium entry per synaptic event (Sobczyk et al., 2005). Moreover, previous studies indicate that GluN2A-containing NMDARs are generally inserted in the PSD whereas GluN2B-containing NMDARs are localized extrasynaptically and associated with SAP102 (Tovar and Westbrook, 1999; Hardingham et al., 2002; Townsend et al., 2003; Washbourne et al., 2004; Hardingham and Bading, 2010). To determine more directly the effects of our biochemical and electrophysiological findings on calcium dynamics, we analyzed calcium signals in WT and *Ppt1^-/-^* neurons transfected with the genetically encoded calcium sensor, GCaMP3 (Tian et al., 2009).

While WT neurons exhibited primarily compartmentalized, dendritic spine-specific calcium signals (**Figure 7A-C**, left, see Video 1), *Ppt1^-/-^* neurons demonstrated diffuse calcium influxes that spread through the dendritic shaft (**Figure 7A-C**, right, see Video 2). These extrasynaptic transients appear rarely in WT cells (**Figure 7**, see Videos). To analyze the calcium dynamics in more detail, measurements of ΔF/F_0_ were made for each dendritic segment, from each cell over the course of the captured videos (see Methods). Multiple transients from the same synaptic site are shown as a heat map of ΔF/F_0_ measurements and they are largely consistent across time in both WT and *Ppt1^-/-^* neurons (**Figure 7B**). Further, plotting of the averaged ΔF/F_0_ transients at an individual synaptic site demonstrates that local fluorescence increases in WT cells are confined to a short distance from the peak ΔF/F_0_ at synaptic sites (**Figure 7B** and C, left), while those of *Ppt1^-/-^* neurons diffuse longer distances within the dendrite (**Figure 7B** and C, right). To quantitatively compare these properties, we performed measurements of area under the curve (AUC) and calcium diffusion distance (see shaded region in **Figure 7C**) for each synaptic site from WT and *Ppt1^-/-^* neurons. These analyses revealed a robust increase in both the AUC (**Figure 7D**) and the calcium diffusion distance (**Figure 7E**) in *Ppt1^-/-^* neurons compared to WT. Furthermore, performing correlation analysis of calcium events across time (see Methods) within a given neuron demonstrates that calcium influxes are more synchronous (increased correlation coefficient) in *Ppt1^-/-^* neurons compared to WT (**Figure 7F**). This result may involve the mechanisms underlying synaptic cluster plasticity, including synaptic integration via translational activation (SITA) influenced by excessive Ca^2^+ entry in *Ppt1^-/-^* neurons (Govindarajan et al., 2006). Together, these data indicate that calcium entry and dispersion are enhanced at *Ppt1^-/-^* synapses *in vitro*.

**Figure 7.**
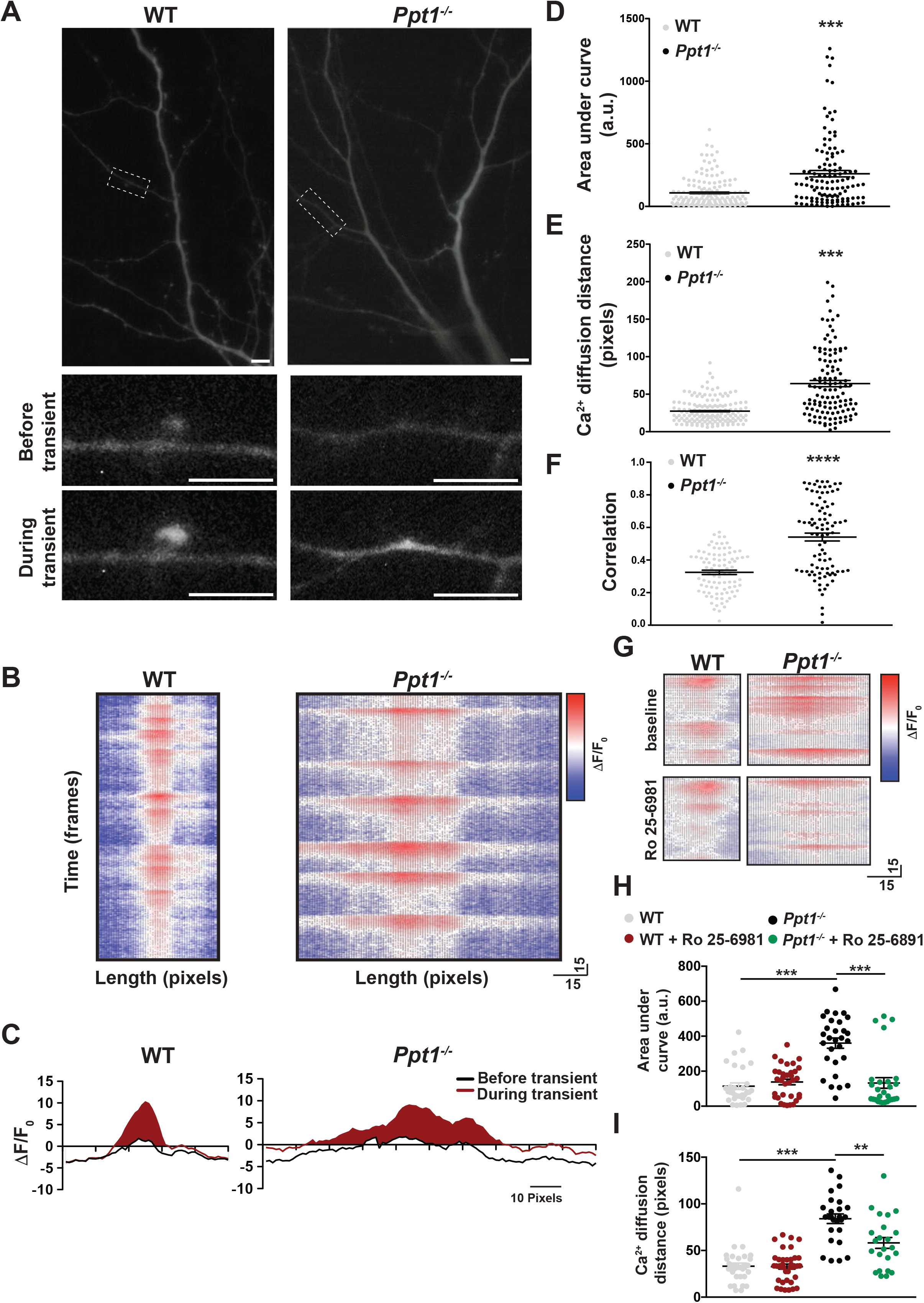
Calcium imaging reveals extrasynaptic calcium dynamics in *Ppt1^-/-^* neurons. DIV16-18, WT and *Ppt1^-/-^* cortical neurons transfected with GCaMP3 and imaged in the absence of Mg^2^+ for 5 minutes. **(A)** Single frames from Videos 1 and 2 of WT (left, note that cell is rotated 90° from Video 1) and *Ppt1^-/-^* (right) cultured neurons. Dendritic segments within the dotted-lines represent zoomed-in images of a single spine (left, WT) or dendritic shaft segment (right, *Ppt1^-/-^*) at baseline (top) and active (bottom) states. Scale=10μm. **(B)** Representative heat maps of ΔF/F_0_ values at one synaptic site from WT (left) and *Ppt1^-/-^* (right) dendrite segments during a portion the imaging session (350 frames, 50 seconds). **(C)** Representative averaged ΔF/F_0_ responses at one synaptic site from WT (left) and *Ppt1^-/-^* (right) neurons. Area under the curve represents calcium influx and is shaded in red. **(D)** Quantification of calcium transient area under the curve WT and *Ppt1^-/-^* neurons. **(E)** Quantification of calcium transient diffusion distance from WT and *Ppt1^-/-^* neurons. **(F)** Quantification of average correlation coefficient (synaptic synchrony) across time between sites of synaptic activity in WT and *Ppt1^-/-^* neurons. **(G)** Representative heat maps of ΔF/F_0_ values at one synaptic site from WT (left) and *Ppt1^-/-^* (right) dendrite segments before (top) and after (bottom) treatment with Ro 25-6981 (130 frames, 18 seconds). **(H)** Quantification of calcium transient area under the curve WT and *Ppt1^-/-^* neurons before and after treatment with Ro 25-6981. **(I)** Quantification of calcium transient diffusion distance from WT and *Ppt1^-/-^* neurons before and after treatment with Ro 25-6981. For experiments in **Figure 7D-E**, *Ppt1^-/-^* and WT were compared (n=170 synaptic sites **(WT)**, n=131 synaptic sites (*Ppt1^-/-^*), 3-6 neurons/group, 3 individual experiments) by t-test and the significance was indicated as follows: ***p<0.001 vs. WT by t-test. For experiments in **Figure 7F**, *Ppt1^-/-^* and WT were compared (n=100 synaptic sites (WT), n=100 synaptic sites (*Ppt1^-/-^*); 3 neurons/group, 2 individual experiments) by t-test and the significance was indicated as follows: ***p<0.001 vs. WT by t-test. For experiments in **Figure 7G-H**, *Ppt1^-/-^* and WT were compared (n=25 synaptic sites (WT), n=28 synaptic sites (*Ppt1^-/-^*); 3 neurons/group, 2 individual experiments) by t-test and the significance was indicated as follows: ***p<0.001 vs. WT by t-test. 65 pixels is representative of 10μm. Error bars represent s.e.m.

These data are in line with our biochemical and electrophysiological findings and suggest that GluN2B-containing NMDARs mediate the observed calcium signals. To further test this possibility, we next treated WT and *Ppt1^-/-^* neurons with Ro 25-6981 (1μM, added in imaging medium following 2.5min imaging at baseline), a GluN2B-containing NMDAR specific antagonist, and performed calcium imaging. Ro 25-6981 had virtually no effect on calcium signals recorded from WT cells (**Figure 7G-I**, see Video 3). In contrast, *Ppt1^-/-^* neurons treated with Ro 25-6981 showed a reduction in dendritic calcium influxes within shafts, while few residual, compartmentalized transients persisted (**Figure 7G-I**, see Video 4). Quantitatively, both AUC (**Figure 7H**) and calcium diffusion (**Figure 7I**) distance were rescued to WT levels following Ro 25-6981 treatment of *Ppt1^-/-^* neurons. Together, these data suggest that *Ppt1^-/-^* neurons have extrasynaptic calcium signaling compared to WT that is sensitive to GluN2B-NMDAR blockade.

### *Ppt1^-/-^* cultured neurons show enhanced vulnerability to NMDA-mediated excitotoxicity

GluN2B-predominant NMDARs are implicated in enhanced neuronal susceptibility to NMDA-mediated neuronal death (Hardingham and Bading, 2002, 2010; Hardingham et al., 2002; Martel et al., 2012). Our results from biochemical, electrophysiological, and live-imaging analyses indicate decreased GluN2A/2B ratio suggesting an intriguing possibility that *Ppt1^-/-^* neurons are more vulnerable to excitotoxicity (Finn et al., 2012). Therefore, we treated WT and *Ppt1^-/-^* cultured neurons with NMDA (varying doses, 10-300μm) and glycine (1-30μm, always in 1:10 ratio with NMDA) for 2 hours and assayed cell viability 24 hours later using the PrestoBlue^®^ reagent (ThermoFisher Scientific) (**Figure 8A**). As expected, both WT and *Ppt1^-/-^* neurons demonstrated dose-dependent reductions in cell viability in response to increasing concentrations of NMDA/glycine (**Figure 8B**). Importantly, *Ppt1^-/-^* neurons were more vulnerable to NMDA insult, as exposure to 10μM NMDA was sufficient to reduce cell viability significantly in *Ppt1^-/-^* neurons but not WT cells (WT = 93 ± 4.1%; *Ppt1^-/-^* = 76 ± 3.5%; \**p*= 0.046; **Figure 8B**). Further, at 100μM NMDA, WT neuron viability decreased by 35%, while *Ppt1^-/-^* neuron viability was reduced significantly further, by 58% (WT = 65 ± 1.8%; *Ppt1^-/-^* = 42 ± 4.5%; \*\**p*= 0.0043; **Figure 8B**). At 300μM NMDA treatment this effect plateaued, as cell viability between WT and *Ppt1^-/-^* neurons was comparable (**Figure 8B**).

**Figure 8.**
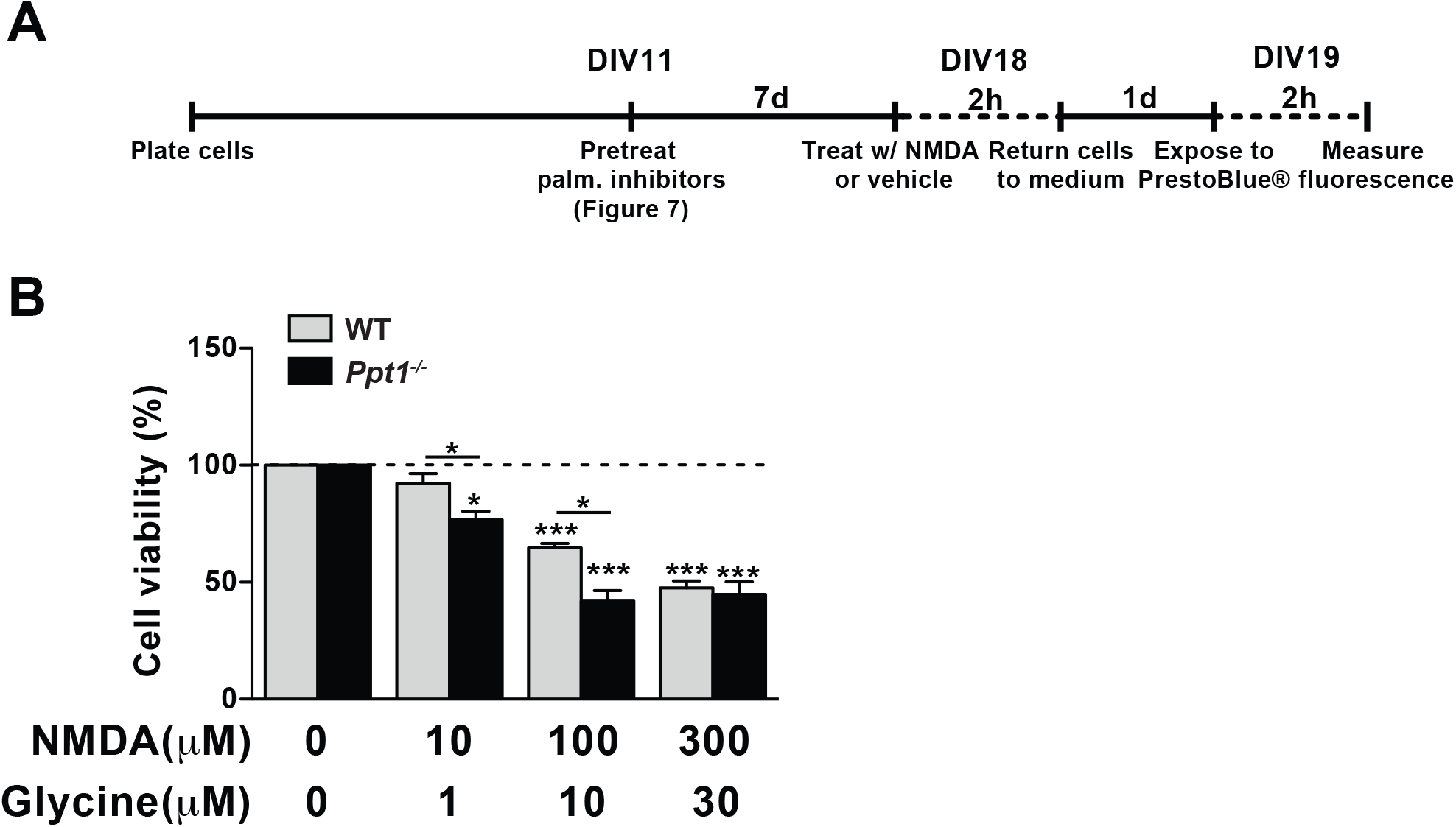
*Ppt1^-/-^* cultured neurons show enhanced vulnerability to NMDA-mediated excitotoxicity. **(A)** Schematic of cellular toxicity experimental design. Briefly, neurons were grown to DIV11, treated with vehicle of palmitoylation inhibitors for 7d (every 48h) and neuronal viability was measured by PrestoBlue^®^ cellular viability assay following exposure (2h exposure, 22h incubation in medium) to NMDA and glycine. **(B)** Quantification of cellular viability in WT and *Ppt1^-/-^* neurons at DIV19 treated with increasing concentrations of NMDA and glycine (10/1, 100/10, and 300/30μm). *Ppt1^-/-^* and WT were compared (n=4 independent experiments, in duplicate) by t-test and significance was indicated as follows: *p<0.05 and ***p<0.001 where indicated; black colored symbols indicate significant differences between vehicle and NMDA/glycine treatment within the same genotype; green colored symbols indicate significant differences between *Ppt1^-/-^* and WT treated with the same concentration of NMDA/glycine. Error bars represent s.e.m.

### Palmitoylation inhibitors rescue enhanced vulnerability to NMDA-mediated excitotoxicity in *Ppt1^-/-^* cultured neurons

We next asked whether this enhanced vulnerability to excitotoxicity is due to hyperpalmitoylation of neuronal substrates, and if it can be corrected by balancing the level of synaptic protein palmitoylation/depalmitoylation. First, we found that 77% of cultured *Ppt1^-/-^* neurons accumulate ALs spontaneously at DIV18-20 (**Figure 9A** and B). Immunostaining for a marker of lysosomal compartments, lysosomal-associated membrane protein-2 (LAMP-2), corroborated these findings, revealing that the observed AL accumulates within lysosomes of *Ppt1^-/-^* but not WT neurons (vehicle treatment in **Figure 9B**). Further, lysosomes appeared swollen in vehicle-treated *Ppt1^-/-^* neurons (see arrows in **Figure 9B**). Treatment with the palmitoylation inhibitors, 2-bromopalmitate (2-BP, 1μM, 7-day treatment) and cerulenin (1μM, 7-day treatment) reduced the percentage of AL-positive neurons (**Figure 9C**) and the area occupied with ALs per neuron (**Figure 9D**). Further, the mean lysosomal size also normalized in *Ppt1^-/-^* neurons when these cells were treated with 2-BP or cerulenin (**Figure 9E**).

**Figure 9.**
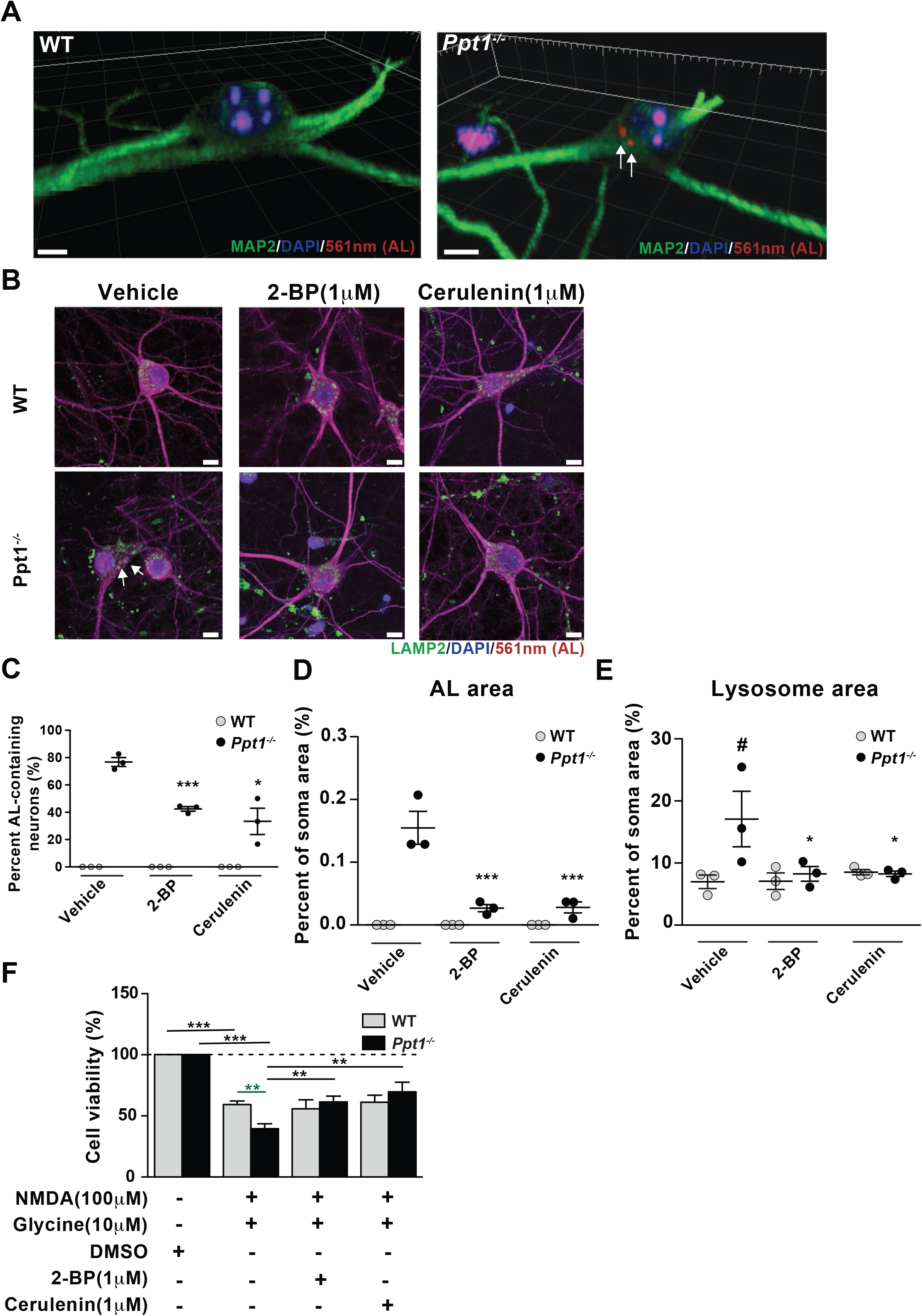
Palmitoylation inhibitors rescue enhanced vulnerability to NMDA-mediated excitotoxicity in *Ppt1^-/-^* cultured neurons. **(A)** 3D reconstructions of a WT and *Ppt1^-/-^* neuron at DIV20. Arrows point to AL deposits. Scale bar=5μm **(B)** Representative collapsed z-stacks of WT and *Ppt1^-/-^* DIV20 neurons, demonstrating accumulations of ALs (arrows) within the soma, particularly within LAMP2-positive vesicles, of *Ppt1^-/-^* neurons. Note the enlarged lysosomes in *Ppt1^-/-^*, vehicle-treated neurons. Scale bar=10μm **(C)** Quantification of the percentage of AL-containing neurons at DIV20 with or without the palmitoylation inhibitors, 2-bromopalmitate (2-BP, 1μM) and cerulenin (1μM), treatment for 6d. **(D)** The percentage of soma area occupied by ALs with or without the palmitoylation inhibitors, 2-BP (1μM) and cerulenin (1μm), treatment for 6d. **(E)** Quantification of the percentage of soma area occupied by lysosomes (LAMP-2-positive vesicles) with and without palmitoylation inhibitor, 2-BP (1μM) and cerulenin (1μM), treatment for 6d. *Ppt1^-/-^* and WT were compared (7-10 neurons/group/experiment, n=3 independent experiments) by t-test and significance was indicated as follows: *p<0.05, ***p<0.001 vs. vehicle-treated *Ppt1^-/-^* neurons; #p<0.05 vs. vehicle-treated WT neurons. **(F)** Quantification of cellular viability in DIV18-20 WT and *Ppt1^-/-^* neurons treated with NMDA and glycine (100/10μM) with or without pretreatment with vehicle (DMSO), 2-BP (1 μM) or cerulenin (1 μM). *Ppt1^-/-^* and WT were compared (n=4 independent experiments, in duplicate) by t-test and significance was indicated as follows: **p<0.01, ***p<0.001 where indicated; black colored symbols indicate significant differences between treatment groups within the same genotype; green colored symbols indicate significant differences between *Ppt1^-/-^* and WT treated with the same concentration of NMDA/glycine. Error bars represent s.e.m.

To examine the efficacy of these compounds in preventing NMDA-mediated toxicity, we pretreated a subset of neurons with the same palmitoylation inhibitors, 2-BP (1μM, DIV12-18) and cerulenin (1μM, DIV12-18) prior to treatment with NMDA and glycine. Notably, pretreatment with both 2-BP and cerulenin improved cell viability of *Ppt1^-/-^* neurons to that of WT following excitotoxicity induction (**Figure 8F**). These results indicate *Ppt1^-/-^* neurons are more vulnerable to excitotoxicity and are consistent with our calcium imaging data that demonstrated the predominance of extrasynaptic, GluN2B-mediated NMDAR activity.

### Palmitoylation inhibitors rescue GluN2B and Fyn kinase hyperpalmitoylation in *Ppt1^-/-^* neurons

Finally, we directly examined the palmitoylation state of neuronal proteins to determine the mechanisms by which hyperpalmitoylation of neuronal substrates may lead to NMDA-mediated excitotoxicity and asked whether palmitoylation inhibitors correct these abnormalities. In particular, we focused on GluN2B palmitoylation, given our evidence implicating GluN2B in the synaptic dysfunction present in *Ppt1^-/-^* neurons. We employed a modified acyl-biotin exchange procedure (Drisdel and Green, 2004), termed the APEGS assay (acyl-PEGyl exchange gel-shift) (Yokoi et al., 2016). The APEGS assay effectively tags the palmitoylation sites of neuronal substrates with a 5kDa polyethylene glycol (PEG) polymer, causing a molecular weight-dependent gel shift in immunoblot analyses. Thus, we quantitatively analyzed the palmitoylated fraction of synaptic proteins and palmitoylated signaling molecules that may influence NMDAR function.

We subjected DIV18 WT and *Ppt1^-/-^* primary cortical neuron lysates to the APEGS assay to determine the palmitoylation state of GluN2B. Indeed, GluN2B palmitoylation was increased in *Ppt1^-/-^* neurons compared to WT at DIV18 (**Figure 10A**). This suggests enhanced surface retention of GluN2B-containing NMDARs in *Ppt1^-/-^* neurons (Mattison et al., 2012). Next, we asked whether palmitoylation inhibitors mitigated excitotoxic vulnerability in *Ppt1^-/-^* neurons by correcting GluN2B hyperpalmitoylation. Both 2-BP and cerulenin treatment (1μm, DIV12-18) as in **Figure 9** normalized levels of GluN2B palmitoylation to those of WT (**Figure 10B**), implying enhanced turnover or reduced surface retention of GluN2B. We also examined another palmitoylated protein, Fyn kinase, as a candidate for facilitating the enhanced surface retention of GluN2B-containing NMDARs. Fyn is a prominent member of the Src kinase family known to directly interact with and affect the synaptic stabilization of GluN2B (Prybylowski et al., 2005; Trepanier et al., 2012). Fyn palmitoylation was increased in *Ppt1^-/-^* neurons compared to WT (**Figure 10C**). Importantly, the palmitoylation of Fyn significantly decreased in both WT and *Ppt1^-/-^* neurons following treatment with 2-BP and cerulenin (**Figure 10D**). Together, these data point to two potentially overlapping mechanisms by which palmitoylation inhibitors reduce the stabilization or retention of GluN2B at the synaptic compartment, thereby reducing cellular calcium load in *Ppt1^-/-^* neurons and mitigating the enhanced susceptibility to excitotoxicity. Further, these data implicate the palmitoylation of Fyn kinase, a molecule that is being targeted for the treatment of Alzheimer’s disease (Nygaard et al., 2014, 2015; Kaufman et al., 2015), in the progression of CLN1.

**Figure 10.**
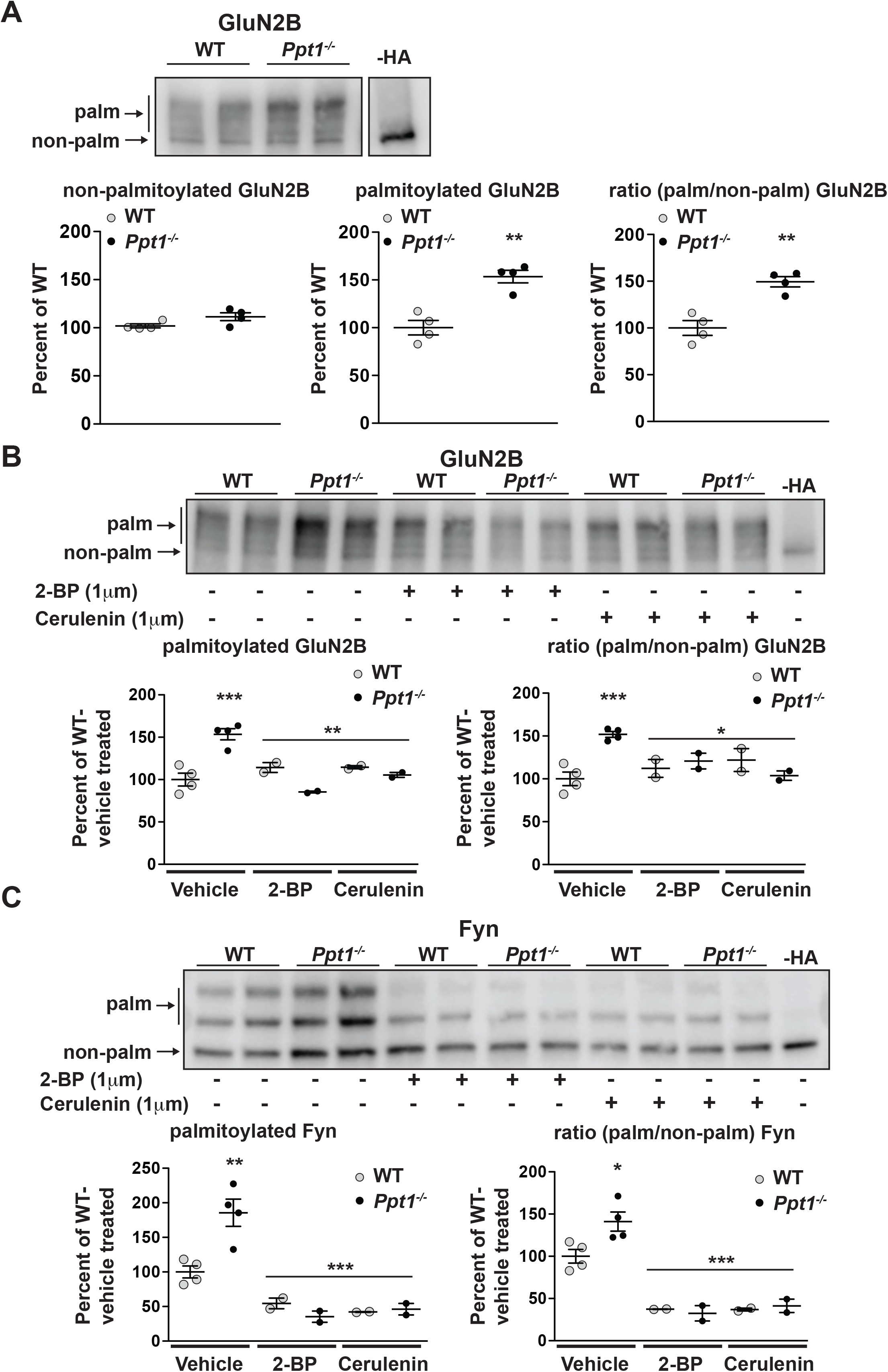
Hyperpalmitoylation of GluN2B and Fyn kinase is reversed in *Ppt1^-/-^* primary cortical neurons by palmitoylation inhibitor treatment. **(A)** Representative post-APEGS immunoblot with minus hydroxylamine (-HA) control (top), and quantification of non-palmitoylated (left), palmitoylated (middle), and the ratio of (palmitoylated/non-palmitoylated, right) GluN2B levels at DIV18 (bottom). **(B)** Representative post-APEGS immunoblot with minus hydroxylamine (-HA) control (top), and quantification of palmitoylated (left), and the ratio of (palmitoylated/non-palmitoylated, right) GluN2B levels following chronic (7d) treatment with the palmitoylation inhibitors, 2-BP (1μm) or cerulenin (1μm) or vehicle control where indicated (bottom). **(C)** Representative post-APEGS immunoblot with minus hydroxylamine (-HA) control (top), and quantification of palmitoylated (left), and the ratio of (palmitoylated/non-palmitoylated, right) Fyn levels following chronic (7d) treatment with the palmitoylation inhibitors, 2-BP (1μm) or cerulenin (1μm) or vehicle control where indicated (bottom). For experiments in **Figure 10A**, *Ppt1^-/-^* and WT were compared (n=4 independent experiments) by t-test and significance was indicated as follows: **p<0.01. For experiments in **Figure 10B-C**, *Ppt1^-/-^* and WT were compared (n=2-4 independent experiments) at each time point using two-way ANOVA followed by Bonferroni post hoc test and the significance was indicated as follows: *p<0.05, **p<0.01 ***p<0.001. Error bars represent s.e.m.

## Discussion

The first mouse model of CLN1 was developed in 2001 by Gupta and colleagues (Gupta et al., 2001). Since the development of this, and alternative models of CLN1, much progress has been made in understanding the temporal, regional, and cell-type specific effects of lipofuscin accumulation and neuronal degeneration, particularly in late stage disease (Gupta et al., 2001; Bible et al., 2004; Kielar et al., 2007; Bouchelion et al., 2014). In addition, comprehensive data characterizing the behavioral dysfunction of the *Ppt1^-/-^* mouse has recapitulated clinical symptoms of the disease (Dearborn et al., 2015). Recent data demonstrate that PPT1 localizes to synaptic compartments and influences presynaptic localization and mobility of prominent presynaptic proteins, including SNAP25 (Kim et al., 2008). These findings correlate with histological and electrophysiological findings in cultured *Ppt1^-/-^* neurons, demonstrating a depletion of presynaptic vesicle pool size (Virmani et al., 2005). Moreover, presynaptic protein localization and function are altered in CLN1 models and in human tissue (Kanaani et al., 2004; Virmani et al., 2005; Kim et al., 2008; Aby et al., 2013).

In the current study, we examined the role of PPT1 in cortical development and postsynaptic maturation in the *Ppt1^-/-^* mouse model of CLN1. Our principal finding demonstrates a role for PPT1 and, more broadly, protein depalmitoylation in the regulation of NMDAR composition and function during visual cortical development. Specifically, we show reductions of GluN2A-containing NMDARs and their preferential synaptic scaffold, PSD-95, in the *Ppt1^-/-^* mouse visual cortex at distinct developmental time points (P33-P60). These biochemical alterations are recapitulated *in vitro* and correlate with immature dendritic spine morphology *in vivo* and *in vitro*, extrasynaptic calcium transients, and enhanced susceptibility to NMDA-induced excitotoxicity. These data are in agreement with previous studies demonstrating markedly enhanced NMDA-, but not AMPA-, mediated toxicity in *Ppt1^-/-^* neurons and improved behavioral phenotype of *Ppt1^-/-^* mice treated with the NMDAR antagonist, memantine (Finn et al., 2012, 2013). Furthermore, *Ppt1^-/-^* neurons exhibit hyperpalmitoylation of both GluN2B and Fyn kinase, which facilitates retention of GluN2B-containing receptors on the cell surface. This dysregulation likely deviates the GluN2A/2B composition and spine morphology toward immaturity, causing enhanced vulnerability to excitotoxic insult. Importantly, chronic palmitoylation inhibitor treatment alleviates *Ppt1^-/-^*-induced dysfunction. Together, these data implicate dysregulated GluN2 subunit switch as a major pathogenic mechanism in CLN1.

### GluN2B to GluN2A subunit switch

Numerous studies have demonstrated the biochemical and functional switching of GluN2B-to GluN2A-containing NMDARs in various brain regions, including the cortex, hippocampus, cerebellum, and brainstem (Paoletti et al., 2013). This developmental event is experience-dependent. Specifically, acute *ex vivo* hippocampal preparations from young (P2-P9) rats reveal rapid, evoked activity-dependent synaptic GluN2A incorporation in response to LTP-induction (Bellone and Nicoll, 2007). Further. Dark rearing rats, effectively removing visual stimuli and consequent afferent activity, delays the GluN2 subunit switch and extends critical period plasticity (Carmignoto and Vicini, 1992; Quinlan et al., 1999b; Philpot et al., 2001). Conversely, environmental enrichment causes an acceleration of visual cortical maturation (Cancedda et al., 2004; Sale et al., 2004; Guzzetta et al., 2009). Here, we have identified the novel mechanism whereby impaired GluN2B to GluN2A subunit switch contributes to the core pathophysiology of a pediatric neurodegenerative disease.

We have demonstrated a novel role for PPT1 in the regulation of palmitoylated postsynaptic proteins. In particular, lack of PPT1 disrupts the NMDA receptor GluN2B to GluN2A subunit switch characteristic of excitatory synaptic maturation. Initially, we predicted synaptic markers, particularly PSD-95, would be overrepresented at postsynaptic sites, since their synaptic distribution depends on the balance between palmitoylation and depalmitoylation (Craven et al., 1999; El-Husseini et al., 2000; Jeyifous et al., 2016). As PSD-95 facilitates the GluN2 subunit switch and preferentially interacts with GluN2A, we also hypothesized an increase in the GluN2A subunit. However, our biochemical data indicate reductions in the total amount as well as the synaptic incorporation of GluN2A and PSD-95 in the PPT1-deficient brain (**Figure 2**, **Supplementary Figure 1**). Interestingly, a recent study suggests that PSD-95 is depalmitoylated by ABHD17 family enzymes, not PPT1 (Yokoi et al., 2016). Further, the palmitoylation state of PSD-95 is reduced with increased synaptic activity (El-Husseini et al., 2002). Consequently, we now postulate that the lack of functional PPT1 results in surface retention of GluN2B-containing NMDAR complexes, either directly or indirectly via the mechanisms discussed below, thereby impeding the developmental switch to GluN2A-containing receptors.

It remains unclear whether impaired GluN2B to GluN2A subunit switch in *Ppt1^-/-^* neurons is entirely attributable to GluN2B hyperpalmitoylation or if Fyn hyperpalmitoylation is also involved. Fyn kinase is developmentally regulated (Umemori et al., 1992; Inomata et al., 1994) and a major palmitoylated downstream kinase of reelin signaling that affects GluN2B surface stabilization (Alland et al., 1994; Koegl et al., 1994; Prybylowski et al., 2005; Kang et al., 2008). Moreover, Fyn kinase is currently being investigated in phase 3 clinical trials as a pharmacological target in AD (Nygaard et al., 2014, 2015; Kaufman et al., 2015). Thus, Fyn dysfunction (in varying contexts) may be a common feature of these neurodegenerative phenotypes. While protein palmitoylation is associated with many of the signaling pathways underlying synaptic development, further study is needed to elucidate precisely (i.e. the direct substrates) how PPT1 influences the GluN2B to GluN2A switch and if Fyn is indeed a key mediator. Studying developmental mechanisms that are 1) associated with GluN2 switching *in vivo* and 2) regulated by protein palmitoylation/depalmitoylation will likely reveal targets for pharmacological intervention in CLN1.

Several other mechanisms underlying this GluN2 subunit switch have been proposed, including reelin, Wnt-5a, and mGluR5 signaling (Groc et al., 2007; Cerpa et al., 2011; Matta et al., 2011). The accumulation of reelin at excitatory synapses during development, for example, mobilizes GluN2B-containing NMDARs and enhances the synaptic contribution of GluN2A-containing NMDARs (Groc et al., 2007; Iafrati et al., 2013). Similarly, evoked activation of mGluR5 at hippocampal synapses is necessary for incorporation of GluN2A-containing NMDARs, and mGluR5-null mice demonstrate deficient GluN2B to GluN2A switching (Matta et al., 2011). Importantly, Wnt-5a and mGluR5 function are directly regulated by palmitoylation (Kurayoshi et al., 2007; Yokoi et al., 2016), suggesting that disruptions in protein depalmitoylation may lead to impaired synaptic maturation through several pathways.

The current work supports a role for PPT1 in regulating postsynaptic proteins. However, whether the pre- or postsynaptic site is affected foremost in *Ppt1^-/-^* mouse visual cortex during development is still unknown. Indeed, we detect a significant reduction in GluN2A and PSD-95 beginning at P33, but not before, while lipofuscin deposition begins as early as P14. However, our electrophysiological, biochemical, and *in vitro* calcium imaging data suggest that this reduction is manifested postsynaptically. Furthermore, we show direct evidence for hyperpalmitoylation of two postsynaptic proteins, the GluN2B subunit and Fyn. One direction of our future work is to examine in more detail the timing, subcellular specificity, and trafficking of both pre- and post-synaptic proteins in *Ppt1^-/-^* mouse.

### Excitotoxicity and NMDAR regulation

Patients afflicted with later-onset NCLs typically exhibit an enlarged VEP prior to degeneration, concurrent with seizure (Pampiglione and Harden, 1977; Pagon et al., 1993; Haltia, 2006). While this phenomenon has not been directly observed in CLN1, it is plausible that disrupted GluN2 subunit switch contributes to hyperexcitability and accelerates cell death leading to the rapid degeneration of neuronal circuits before the diagnosis can be made. In line with this notion, NMDAR subunit dysregulation is implicated in various neuropsychiatric and neurodegenerative disorders (Paoletti et al., 2013; Zhou and Sheng, 2013). Furthermore, recent evidence describing the effects of GluN2 subunit incorporation on NMDAR function suggests a mechanism linking subunit specificity to excitotoxicity and neuronal degeneration (Martel et al., 2009, 2012; Hardingham and Bading, 2010). GluN2 subunit composition and NMDAR localization activate opposing downstream transcriptional programs. Specifically, GluN2A-containing NMDARs in the postsynaptic density activate cyclic-AMP response element binding protein (CREB) and other transcription factors associated with cell-survival and learning. In contrast, activation of GluN2B-containing, extrasynaptic NMDARs preferentially triggers pro-apoptotic signaling pathways and causes inhibition of CREB (Hardingham and Bading, 2002; Hardingham et al., 2002). Though this system is likely more intricate than described here (Thomas et al., 2006), these previous studies are consistent with our observations that *Ppt1^-/-^* neurons are biased toward extrasynaptic calcium transients (**Figure 7** and Video 2) and that they are more susceptible to excitotoxicity (**Figure 8**). Furthermore, the most significant outcome of this study is that palmitoylation inhibitors mitigated the pro-apoptotic predisposition of *Ppt1^-/-^* neurons *in vitro* (**Figure 9**).

The incorporation of GluN2A into NMDARs is experience-dependent (Stocca and Vicini, 1998; Quinlan et al., 1999a, 1999b). An intriguing possibility is that *Ppt1^-/-^* neurons in sensory cortices are unable to tolerate normal sensory experiences, in part because this experience-dependent GluN2 subunit switch is disrupted. Indeed, PPT1-defeciency results in selective degeneration of thalamic nuclei and primary sensory cortices (Bible et al., 2004; Kielar et al., 2007). Further, PPT1 expression is developmentally-regulated in WT rodents, such that functional PPT1 increases with cortical maturation and peaks in early adulthood, when it may regulate this switching phenomenon (Suopanki et al., 1999a, 1999b). Together, we argue that intact PPT1 plays a critical role in regulating NMDAR functional properties in response to external stimuli, thereby facilitating synaptic maturation and preventing excitotoxicity. Whether manipulating neuronal activity or experience-dependent synaptic plasticity ameliorates disease progression remains unknown and is a focus of ongoing experiments.

### Implications for other neurodegenerative diseases

While substantial progress has been made in our understanding of adult-onset neurodegenerative diseases including Alzheimer’s disease and Parkinson’s disease, effective, disease-modifying therapeutics are yet to be developed for most of these disorders. In part, this is likely due to the genetic complexity and heterogeneity of these diseases as well as lifestyle and environmental factors limiting the translational success of seemingly promising therapeutic strategies. Recently, studies in monogenic diseases have attracted attention because they share common pathological hallmarks with adult-onset neurodegenerative diseases. This approach has turned out to be valuable to decipher underlying disease mechanisms (Peltonen et al., 2006). For instance, heterozygous mutations in the glucocerebrosidase (*GBA*) gene, which cause Gaucher disease if homozygous, confer risk to Parkinson’s disease (Neudorfer et al., 1996; Tayebi et al., 2001; Sidransky, 2012; Sidransky and Lopez, 2012). The link between the two diseases exemplifies the critical role of lysosomal degradation in inclusion body formation and corroborates mounting evidence for autophagy as a key mechanism to clear neuronal waste and maintain cellular health (Nixon, 2005, 2013).

Lipofuscin is not only the cardinal hallmark of the NCLs, but also accumulates in many neurodegenerative disorders, including Alzheimer’s and Huntington’s diseases. However, it remains inconclusive whether lipofuscin aggregation is an adaptive, neuroprotective mechanism, or a direct cause of neuronal degeneration. Clinical therapies in CLN1 patients aimed at targeting storage material are largely unsuccessful (Gavin et al., 2013; Levin et al., 2014). For instance, clinical trials using small compounds that effectively depleted lipofuscin in CLN1 patients demonstrated minor subjective improvements. Nevertheless, patients progressed to a vegetative state by 52 months (Levin et al., 2014). Therefore, it is likely that CLN1 pathology involves not only lipofuscin deposition, but also an excess of palmitoylated proteins, which alters synaptic functions. Indeed, emerging evidence indicates that both endosomal sorting and lysosomal proteolysis dynamically contribute to mechanisms underlying synaptic plasticity (Shehata et al., 2012; Gokhale et al., 2015; Goo et al., 2017). Despite detectable NMDAR disruption not beginning before P33, it remains plausible that synaptic dysregulation of some type starts either before or concurrent with the deposition of storage material in CLN1 disease. Understanding synaptic mechanisms that are regulated by PPT1 and other NCL-associated gene products will shape the foundation for treating these devastating diseases. Our data point to NMDARs as a potential therapeutic target and corroborate the efficacy of memantine in *Ppt1^-/-^* mice (Finn et al., 2013). However, other receptor subunits and ion channels also undergo palmitoylation, including AMPARs (Hayashi et al., 2005) and GABARs (Fang et al., 2006). Further studies are needed to reveal the role of PPT1 in regulating these palmitoylated synaptic proteins.

Lastly, several proteins associated with adult-onset neurodegenerative disorders, such as amyloid precursor protein (APP) and huntingtin, are palmitoylated and regulate synaptic functions (Huang et al., 2004; Smith et al., 2005; Zheng and Koo, 2006; Bhattacharyya et al., 2013). Therefore, understanding how the balance between protein palmitoylation and depalmitoylation affects neuronal functions has broad implications and will have a novel impact on the therapeutic strategies against these and other brain disorders.

In conclusion, we have demonstrated that an imbalance between protein palmitoylation and depalmitoylation accelerates lipofuscin accumulation. Further, we have shown that lack of PPT1 function results in the stagnation of developmental GluN2 subunit switch, leading to enhanced vulnerability to glutamate-mediated excitotoxicity. Our results indicate a vital role for PPT1 in the regulation of postsynaptic maturation and set the stage for further investigating protein depalmitoylation in NCLs as well as adult-onset neurodegenerative diseases associated with lipofuscin.

## Materials and methods

### Animals

All animal procedures were performed in accordance with the guidelines of the University of Illinois of Chicago Institutional Animal Care and Use Committee. *Ppt1^+/−^* (heterozygous) mice were obtained from Jackson Laboratory and maintained on 12h light/dark cycle with food and water *ad libitum*. Breeding of *Ppt1^+/−^* animals results in litters containing *Ppt1^-/-^, Ppt1^+/−^*, and *Ppt1^+/+^* (WT) animals. *Ppt1^-/-^* and WT littermate controls at specified developmental time points: P11, P14, P28, P33, P42, P60, P78, and P120 were genotyped in-house (Gupta et al., 2001) and used for experiments.

### Brain fractionation and Western blot

For collection of brain for biochemistry (immunoblot), *Ppt1^-/-^* and WT animals were decapitated following isoflurane anesthesia, then the brain was removed, and washed in ice cold PBS. The occipital cortex (visual cortex), hippocampus, and remaining cortex were separately collected on ice. Isolated visual cortices from *Ppt1^-/-^* and WT animals were homogenized in ice-cold synaptosome buffer (320mM sucrose, 1mM EDTA, 4mM HEPES, pH7.4 containing 1x protease inhibitor cocktail (Roche), 1x phosphatase inhibitor cocktail (Roche) and 1mM PMSF) using 30 strokes in a Dounce homogenizer. Aliquots for whole lysate (WL) were stored and the remaining sample was used for synaptosome preparation, performed as previously with slight modification. In brief, WLs were centrifuged at 1,000 x g to remove cellular debris, supernatant was then centrifuged at 12,000 x g for 15min to generate pellet P2. P2 was resuspended in synaptosome buffer and spun at 18,000 x g for 15min to produce synaptosomal membrane fraction, LP1, which was used for downstream biochemical analyses (synaptosomes). For immunoblot, protein concentration of each sample was determined using BCA protein assay (Pierce). Samples were then measured to 20μg total protein in 2x Laemmli buffer containing 10% β-mercaptoethanol (Bio-rad), boiled at 70°C for 10min and loaded into 10% tris-glycine hand cast gels (Bio-rad), or 4-20% precast gels (Bio-rad) for electrophoresis (110V, 1.5-2h). Proteins were wet-transferred to PVDF membranes (Immobilon-P, Millipore), blocked in TBS, pH7.4 containing 5% non-fat milk and 0.1% Tween-20 (TBS-T+5% milk). Membranes were incubated in primary antibody solutions containing 2% BSA in TBS-T for 2h at RT or overnight at 4°C. Primary antibodies were used as follows: GluN2A (Cat: NB300-105, 1:1,000, Novus Biologicals), GluN2B (Cat: 75/097, 1:1,000, Neuromab), GluN1 (Cat: 75/272, 1:1000, Neuromab), PSD-95 (Cat: K28/74, 1:2,000, Neuromab), SAP102 (Cat: N19/2, 1:2,000, Neuromab), Fyn kinase (Cat: 4023, 1:1,000, Cell signaling) and β-actin-HRP (Cat: MA5-15739-HRP, 1:2,000, ThermoFisher). Membranes were then incubated with appropriate secondary, HRP-conjugated antibodies (Jackson ImmunoResearch) at either 1:5,000, 1:10,000, or 1:30,000 (PSD-95 only) for 1h at RT. Visualization and quantification was performed using Pierce SuperSignal ECL substrate and Odyssey-FC chemiluminescent imaging station (LI-COR). Signal density for each synaptic protein was measured using the LI-COR software, Image Studio Lite (version 5.2) and was normalized to the signal density for β-actin loading control for each lane. A total of four independent experiments was performed for both WL and LP1 analyses, with a minimum of two technical replicates for each experiment averaged together.

### Histology and autofluorescent lipopigment quantification

*Ppt1^-/-^* and WT mice were anesthetized using isoflurane and transcardially perfused with ice cold PBS (pH 7.4, ~30ml/mouse) followed by 4% paraformaldehyde (PFA) in PBS (~15ml/mouse). Brains were removed and post-fixed for 48h at 4°C in 4% PFA and transferred to PBS, pH7.4 containing 0.01% sodium azide for storage if necessary. Brains from *Ppt1^-/-^* and WT animals were incubated in 30% sucrose solution for 48h prior to sectioning using Vibratome 1000 in cold PBS. For imaging and quantification of AL, sagittal sections were cut at 100μm. Every third section was mounted on Superfrost Plus microscope slides (VWR) using Vectamount mounting media containing DAPI (Vector Laboratories, cat: H-5000). Interlaced/overlapping images of visual cortex area V1 from the cortical surface to subcortical white matter (or subiculum), which was localized using Paxino’s mouse atlas (sagittal), were collected for 2-4 sections from each animal using a Zeiss LSM710 confocal laser scanning microscope at 40x magnification (excitation at 405nm to visualize DAPI and 561nm to visualize AL). All sections were imaged using identical capture conditions. Quantification of AL was performed by thresholding images in FIJI (NIH), generating a binary mask of AL-positive pixels (satisfied threshold) vs. background. The identical threshold was applied to each image (from cortical surface to subcortical white matter and across animals). Percent area occupied by AL puncta that satisfied the threshold was then calculated using the “analyze particles” tool in FIJI. This analysis was performed for 2-4 sections (total of ~10-20 images, as imaging an entire cortical column is typically 5 interlaced images) from each animal and averaged together to give a single value, representative of the total area occupied by AL in the cortical column imaged. Three to six animals per group were analyzed this way and averaged to give the mean area occupied by AL at each time point, for both genotypes (n=4-6 animals/group).

### Electrophysiology

WT and *Ppt1^-/-^* animals at P42 were deeply anesthetized using isoflurane drop method and decapitated. Brains were resected in semi-frozen oxygenated (95% O2 and 5% CO2) artificial cerebrospinal fluid (aCSF, in mM: NaCl 85, sucrose 75, KCl 2.5, CaCl2 0.5, MgCl_2_ 4, NaHCO_3_ 24, NaH_2_PO_4_ 1.25, D-glucose 25, pH 7.3), and 350μm sections containing visual cortex area V1 were sectioned using a Leica VT1200 S vibratome in semi-frozen aCSF. After recovery (1h) in aCSF at 30°C, sections were transferred to the recording chamber, perfused at 2ml/min with aCSF at 30°C. Following localization of visual cortex area V1 using Paxinos mouse brain atlas, a stimulating electrode was placed in layer IV, and pyramidal neurons from layer II/III were blindly patched (patch solution in mM: CsOH monohydrate 130, D-Gluconic acid 130, EGTA 0.2, MgCl_2_ 1, CsCl 6, Hepes 10, Na_2_-ATP 2.5, Na-GTP 0.5, Phoshocreatine 5, QX-314 3; pH 7.3, osmolarity 305 mOsm) and recorded in voltage clamp mode at +50mV (V_H_) to remove Mg^2^+ block from NMDARs. NMDA-EPSCs were pharmacologically isolated via addition of CNQX (10μM), (+)-Bicuculline (60 μM) and SCH 50911 to block AMPA, GABAa and GABAb receptors, respectively. Stimulation intensity was titrated to give a saturating postsynaptic response, and EPSCs were then recorded, averaging 5-10 sweeps. The decay phase of the averaged NMDAR-EPSCs were then fitted to a double exponential (Carmignoto and Vicini, 1992). We calculated for each cell: the amplitude of the fast (Af) component (GluN2A-mediated), the amplitude of the slow (As) component (GluN2B-mediated), the contribution of the fast component Af/ Af+As to the overall decay phase, the τ fast (τf), the τ slow (τs) and the τ weighted (τw) in WT and *Ppt1^-/-^* mice following this formula: τw= τfx(Af/Af+As) + τsx(As/Af+As) (n=10/4 (cells/animals), WT; n=12/5 PPT-KO).

### *In utero* electroporation

*In utero* electroporation was performed as previous(Yoshii et al., 2011, 2013). Timed-pregnant dams at E16.5 were deeply anesthetized via isoflurane (3% induction, 1-1.5% for maintenance of anesthesia during surgery) and laparotomized. The uterus was then externalized and up to ~1μl of solution containing GFP construct (2μg/μl) and fast green dye was delivered into the left lateral ventricle through the uterine wall using a micropipette. Using an ECM 830 Square Wave electroporator (Harvard Apparatus, Holliston MA), brains were electroporated with 5 pulses of 28V for 50msec at intervals of 950msec at such an angle to transfect neurons in visual cortex. After recovery, pregnancies were monitored and pups were delivered and nursed normally. Electroporated pups were genotyped, raised to P33, and sacrificed via transcardial perfusion as described above. Electroporated brains from WT and *Ppt1^-/-^* mice (procedure schematized in **Figure 4A**) were sectioned and sequentially mounted. Transfected neurons in visual cortex (**Figure 4B**) were imaged to capture all apical neurites and 3D reconstructed images were analyzed in Imaris (Bitplane) for dendritic spine characteristics known to be associated with synaptic maturity (spine length, spine volume, and spine head volume). At least two z-stack images (typically >100 z-planes/image) were stitched together to capture the prominent apical neurites and extensions into the cortical surface for each cell. Each stitched image, equivalent to one cell, was considered one n.

### Primary cortical neuron culture

For primary cortical neuron cultures, embryos from timed-pregnant, *Ppt1^−/+^* dams were removed, decapitated, and cortices resected at embryonic day (E) 15.5. All dissection steps were performed in ice cold HBSS, pH7.4. Following cortical resection, tissue from each individually-genotyped embryo were digested in HBSS containing 20U/ml papain and DNAse (20min total, tubes flicked at 10min) before sequential trituration with 1ml (~15 strokes) and 200μl (~10 strokes) pipettes, generating a single-cell suspension. For live-cell imaging experiments, cells were counted then plated at 150,000-180,000 cells/well in 24-well plates containing poly-D-lysine/laminin-coated coverslips. For biochemical experiments, i.e. immunoblot, APEGS assay *in vitro*, cells were plated on poly-D-lysine/laminin-coated 6-well plates at 1,000,000 cells/well. Cells were plated and stored in plating medium (Neurobasal medium containing B27 supplement, L-glutamine and glutamate) for 3-5 DIV, before replacing half medium every 3 days with feeding medium (plating medium without glutamate). Cultures used in chronic palmitoylation inhibitor treatment were exposed to either DMSO (vehicle), 2-BP (1μm, Sigma, cat: 238422) or cerulenin (1μm, Cayman Chemicals, cat: 10005647) every 48 hours between DIV 11 and 18.

### Primary cortical neuron harvest and immunoblotting

Primary cortical neurons from E15.5 WT and *Ppt1^-/-^* embryos were cultured for 7, 10, or 18 DIV prior to harvest for immunoblot or APEGS assay (only DIV18 used for APEGS). To harvest protein extracts, cells were washed 2x with ice-cold PBS before addition of lysis buffer containing 1% SDS and protease inhibitor cocktail, 500μl/well. Cells were incubated and swirled with lysis buffer for 5 minutes, scraped from the plate, triturated briefly, and collected in 1.5ml tubes. Lysates were centrifuged at 20,000g for 15min to remove debris, and the supernatant was collected for biochemical analysis. Immunoblotting analyses were performed as in section 2.2. APEGS assay was carried out as described in section 2.7.

### APEGS assay on primary cortical neuron lysates

The APEGS assay was performed as utilized in Yokoi, 2016 and recommended by Dr. M. Fukata (personal communication, 06/2018). Briefly, cortical neuron lysates were brought to 150μg total protein in a final volume of 0.5ml buffer A (PBS containing 4% SDS, 5mM EDTA, protease inhibitors, remaining sample used in aliquots for “input”). Proteins were reduced by addition of 25mM Bond-Breaker™ TCEP (0.5M stock solution, ThermoFisher) and incubation at 55°C for 1h. Next, to block free thiols, freshly prepared N-ethylmaleimide (NEM) was added to lysates (to 50mM) and the mixture was rotated end-over-end for 3h at RT. Following 2x chloroform-methanol precipitation (at which point, protein precipitates were often stored overnight at −20°C), lysates were divided into +hydroxylamine (HA) and –HA groups for each sample, which were exposed to 3 volumes of HA-containing buffer (1M HA, to expose palmitolylated cysteine residues) or Tris-buffer control (HA, see **Figure 10**), respectively, for 1h at 37°C. Following chloroform-methanol precipitation, lysates were solubilized and exposed to 10mM TCEP and 20mM mPEG-5k (Laysan Bio Inc., cat# MPEG-MAL-5000-1g) for 1h at RT with shaking (thereby replacing palmitic acid with mPEG-5K on exposed cysteine residues). Following the final chloroform-methanol precipitation, samples were solubilized in a small volume (60μl) of PBS containing 1% SDS and protein concentration was measured by BCA assay (Pierce). Samples were then brought to 10μg protein in laemmli buffer with 2% β-mercaptoethanol for immunoblot analyses as in section 2.2. Quantification of palmitoylated vs. non-palmitoylated protein was carried out as in section 2.2, with the added consideration that palmitoylated protein was taken as the sum of all (typically two-three distinct bands, see **Figure 9**) bands demonstrating the APEGS-dependent molecular weight shift compared to the –HA control lane. Non-palmitoylated protein was quantified from the band size-matched to the –HA control sample. The ratio was taken as the palmitoylated protein divided by non-palmitoylated protein, all divided by β-actin control from the same lane.

### Transfection, dendritic spine and calcium imaging analyses

For analysis of dendritic spine morphology, WT and *Ppt1^-/-^* neurons were transfected between DIV6-8 with GFP using Lipofectamine^®^ 2000 (ThermoFisher) according to manufacturer protocol. Briefly, GFP DNA construct (~2μg/μl, added at ~1μg/well) was mixed with Lipofectamine-containing Neurobasal medium, incubated for 30min to complex DNA-Lipofectamine, equilibrated to 37°C, and added to the cells 250μl/well for 1-1.5h. Following incubation, complete medium was returned to the cells. Neurons were then imaged at DIV15 and DIV20 for dendritic spine morphology using a Zeiss LSM 710 confocal microscope equipped with a heated stage at 63x magnification. GFP-positive neurons were imaged at 0.2μm Z-plane interval (typically 25 Z-planes/image). Three to seven overlapping Z-stacks were stitched to visualize an entire neuron. Z-stack images were collapsed into a single plane and dendritic spines were analyzed using semi-automated image processing software, Imaris (Bitplane). The same dendrite and dendritic spine processing parameters were used for each image. For DIV15: n=4-5 neurons/group, 3-independent experiments, WT=21,514 spines; *Ppt1^-/-^*=18,013 spines. For DIV20: n=3 neurons/group, 2-independent experiments, WT=11,335 spines; *Ppt1^-/-^*=9,958 spines.

To directly image calcium signals in WT and *Ppt1^-/-^* neurons, cells were transfected as above using the construct encoding GCaMP3 (see **Acknowledgments**) at DIV6-8. Cells were grown to DIV18 then imaged at room temperature in Tyrode’s solution (imaging medium, 139mM NaCl, 3mM KCl, 17mM NaHCO3, 12mM glucose, and 3mM CaCl2) for a maximum of 15min using a Mako G-507B camera mounted onto a Leica inverted microscope. Videos were acquired at ~7 frames per second using StreamPix software (NorPix). A maximum of 5min per neuron was recorded (thus, minimum 3 neurons per coverslip were acquired). N=3-6 neurons/group, three independent experiments. For treatment with Ro 25-6981, neurons were imaged at baseline for 2-2.5min before adding Ro 25-6981 (1μM) directly to the imaging medium. Neurons were then imaged for an additional 2.5min.

To analyze the area under the curve (AUC) and width (diffusion distance) of calcium transients, 500-600 frames from the middle of each video (average frame count for whole videos= ~2000 frames) for WT and *Ppt1^-/-^* neurons were analyzed using FIJI (NIH). Dendritic segments, excluding primary dendrites, were traced using a segmented line ROI with pixel width of 50, which reliably encompassed the dendritic segment and accompanying dendritic spines. Next, the following macro derived from the ImageJ forum (http://forum.imagej.net/t/how-to-obtain-xy-values-from-repeated-profile-plot/1398) was run on each individual ROI:

~~~
macro “Stack profile Plot” {
     collectedValues=““;
     ymin = 0;
     ymax = 255;
     saveSettings();
     if (nSlices==1)
       exit(“Stack required”);
     run(“Profile Plot Options…”,
        “width=400 height=200 minimum=“+ymin+” maximum=“+ymax+” fixed”);
     setBatchMode(true);
     stack1 = getImageID;
     stack2 = 0;
     n = nSlices;
     **for** (i=1; i<=n; i++) {
                showProgress(i, n);
                selectImage(stack1);
                setSlice(i);
                run(“Clear Results”);
                   profile = getProfile();
                **for** (j=0; j<profile.length; j++) {
                           collectedValues=collectedValues+profile[j] + “\t”;
                }
                collectedValues=collectedValues+”\n”;
                run(“Plot Profile”);
                run(“Copy”);
                w = getWidth; h = getHeight;
                close();
                **if** (stack2==0) {
                           newImage(“Plots”, “8-bit”, w, h, 1);
                           stack2 = getImageID;
                } **else** {
                              selectImage(stack2);
                              run(“Add Slice”);
                }
                run(“Paste”);
     }
     f = File.open(“C:/”cell#, ROI #”.xls”);
     print(f, collectedValues);
     setSlice(1);
     setBatchMode(false);
     restoreSettings();
       }
~~~

This gives the fluorescence intensity at each pixel along the ROI across the time/frame dimension. The background fluorescence for each ROI was then subtracted by averaging the fluorescence across the ROI in an inactive state (no calcium transients), giving the measure ΔF/F_0_ when examined across time/frame. For each ROI (up to 1265 pixels in length), each calcium transient at individual synaptic sites (dendritic spines and adjacent shafts) was averaged. Those averages were then compiled to give the average transient signal, which was then used to analyze the AUC and calcium diffusion distance (WT=55 ROIs, 170 synaptic sites, 1630 transients; *Ppt1^-/-^*=38 ROIs, 131 synaptic sites, 1281 transients; n=3-6 neurons/group/experiment, 3 repetitions). For Ro 25-6981-treated neurons, the same protocol was followed with the exception that calcium transients at an individual synaptic site were split into “before application” and “after application” groups.

To analyze synaptic synchrony, ΔF/F_0_ measurements for 20 randomly-chosen sites of synaptic activity per neuron were correlated across the time dimension (500 frames of each video). A correlation matrix was generated to determine the average correlation of each synaptic site with all other chosen sites. The average values for each synaptic site, for 5 neurons/group are plotted in **Figure 7**.

### NMDA toxicity assays

To measure cell viability following exposure of WT and *Ppt1^-/-^* neurons to NMDA and glycine, neurons were plated as above and grown to DIV18. For experiments presented in **Figure 6**, feeding medium was removed from neurons, stored at 37°C, and replaced with B27-free Neurobasal medium with or without NMDA/glycine at the following concentrations: 10/1 μM, 100/10μM, or 300/30μM (ratio maintained at 10:1). Cells were incubated for 2h at 37°C in treatment medium. Following incubation, treatment medium was removed and replaced with the original feeding medium. Cells were then incubated an additional 22h before addition of PrestoBlue^®^ cell viability reagent (ThermoFisher). At 24h, fluorescence intensity of each well was measured using a Beckman Coulter DTX 800 Multimode Detector. Cell viability for each treatment condition was calculated and expressed as percentage of vehicle-treated control wells (no pretreatment, no NMDA application). Experiments in **Figure 7** were performed similarly except that cultures were pretreated with either DMSO (vehicle), 2-BP (1μM, Sigma, cat: 238422) or cerulenin (1μM, Cayman Chemicals, cat: 10005647) every 48 hours between DIV 11 and 18.

### AL accumulation *in vitro*, palmitoylation inhibitor treatment, imaging and analysis

WT and *Ppt1^-/-^* neurons were cultured as above. To examine AL deposition, neurons were grown to DIV18-20, fixed in 4% PFA for 10min at RT, and stored in PBS for up to 72 hours prior to immunocytochemistry. To examine AL accumulation alone, cells were immunostained for the microtubule associated protein, MAP2 (Millipore Sigma, cat: AB5622) and mounted in DAPI-containing mounting medium. To assess AL localization, DIV18-DIV20 neurons were immunostained for MAP2 and LAMP-2 (Abcam, cat: ab13524). Neurons were then imaged at random using a Zeiss LSM 710 confocal microscope at 63x magnification. Z-stacks (0.4μm Z-plane interval, 12-22 Z-planes/image) were taken at 512 x 512 pixel density. 7-10 neurons/group for three independent experiments.

To semi-automatically analyze the percentage of AL-containing cells, the cytosolic area covered by AL deposits, and the cytosolic area covered by lysosomes, images immunostained for MAP2 and LAMP-2 were processed in FIJI. Each channel of the image: LAMP-2 (488nm), MAP2 (633nm), DAPI (405nm), AL (561nm) was thresholded separately as to display only the lysosomes, cell soma, the nucleus, and AL deposits, respectively. Thresholds were kept identical between images. Next, the areas of these compartments/deposits were measured using the “analyze particles” tool restricted to an ROI tracing the cell soma. Lysosomes needed to have a circularity of >0.5 to avoid counting small clusters of lysosomes as a single unit (Bandyopadhyay et al., 2014; Grossi et al., 2016). To measure AL deposits, the same approach was used with the additional constraint: AL deposits were required to have a circularity >0.4 and comprise more than 8 adjacent pixels. Cytosolic area was calculated by measuring MAP2 signal area and subtracting the area occupied by DAPI stain.

### Immunocytochemistry

Coverslips were stained in runs so that all experimental and control groups were immunostained simultaneously. Coverslips were washed 3x with TBS, permeabilized for 20min at RT with TBS containing 0.5% Triton X-100, and blocked for 1h at RT in TBS containing 0.1% Triton X-100 and 5% BSA. Then, primary antibody (MAP2 or LAMP-2) at 1:400 dilution was added to coverslips in TBS containing 0.1% Triton X-100 and 1% BSA and incubated for 2h at RT or overnight at 4°C. Following 4X washes with TBS containing 0.1%

Triton X-100, cells were incubated with 1:400 secondary, fluorophore-linked antibody (either Alexa Fluor 488, cats: A-11034, A-11006; or Alexa Fluor 633, ThermoFisher, cat: A-21070) in TBS containing 0.1% Triton x-100 and 1% BSA. These steps are repeated for double immunostained cells. For LAMP-2/MAP2 double immunostaining, saponin was used in place of Triton X-100 at the same concentrations. Coverslips are then mounted on SuperFrost Plus slides in DAPI Vectamount medium.

## Acknowledgments

The authors thank Dr. Froylan Calderon de Anda (Universitätsklinikum Hamburg-Eppendorf) for the GCaMP3 construct used to visualize calcium activity in **Figure 6**. This work is supported by startup funding awarded to A.Y. by the University of Illinois at Chicago, Department of Anatomy and Cell Biology.

## Author contributions statement

A.Y. conceptualized and oversaw experiments, provided resources, prepared and revised the manuscript. K.P.K designed and performed experiments, performed data analysis, prepared and revised the manuscript as well as the figures. W.F. and F.B. performed electrophysiological recordings (**Figure 3**) and aided in the revision of the manuscript. S.A. and R.S. analyzed calcium dynamics data (**Figure 7**) and aided in manuscript revision.

---

**Video 1. Spontaneous calcium activity in DIV16-18 WT neuron.** Representative video of spontaneous neuronal calcium activity in a WT cultured neuron at DIV16-18.

**Video 2. Spontaneous calcium activity in DIV16-18 *Ppt1^-/-^* neuron.** Representative video of spontaneous neuronal calcium activity in a *Ppt1^-/-^* cultured neuron at DIV16-18.

**Video 3. Spontaneous calcium activity in DIV16-18 WT neuron before and after treatment with Ro 25-6981.** Representative video of spontaneous neuronal calcium activity in a WT cultured neuron at DIV16-18 prior to, and following, bath application of Ro 25-6981 (1μM, after 30 seconds).

**Video 4. Spontaneous calcium activity in DIV16-18 *Ppt1^-/-^* neuron before and after treatment with Ro 25-6981.** Representative video of spontaneous neuronal calcium activity in a *Ppt1^-/-^* cultured neuron at DIV16-18 prior to, and following, bath application of Ro 25-6981 (1μM, after 31 seconds).

**Supplementary Table 1.**
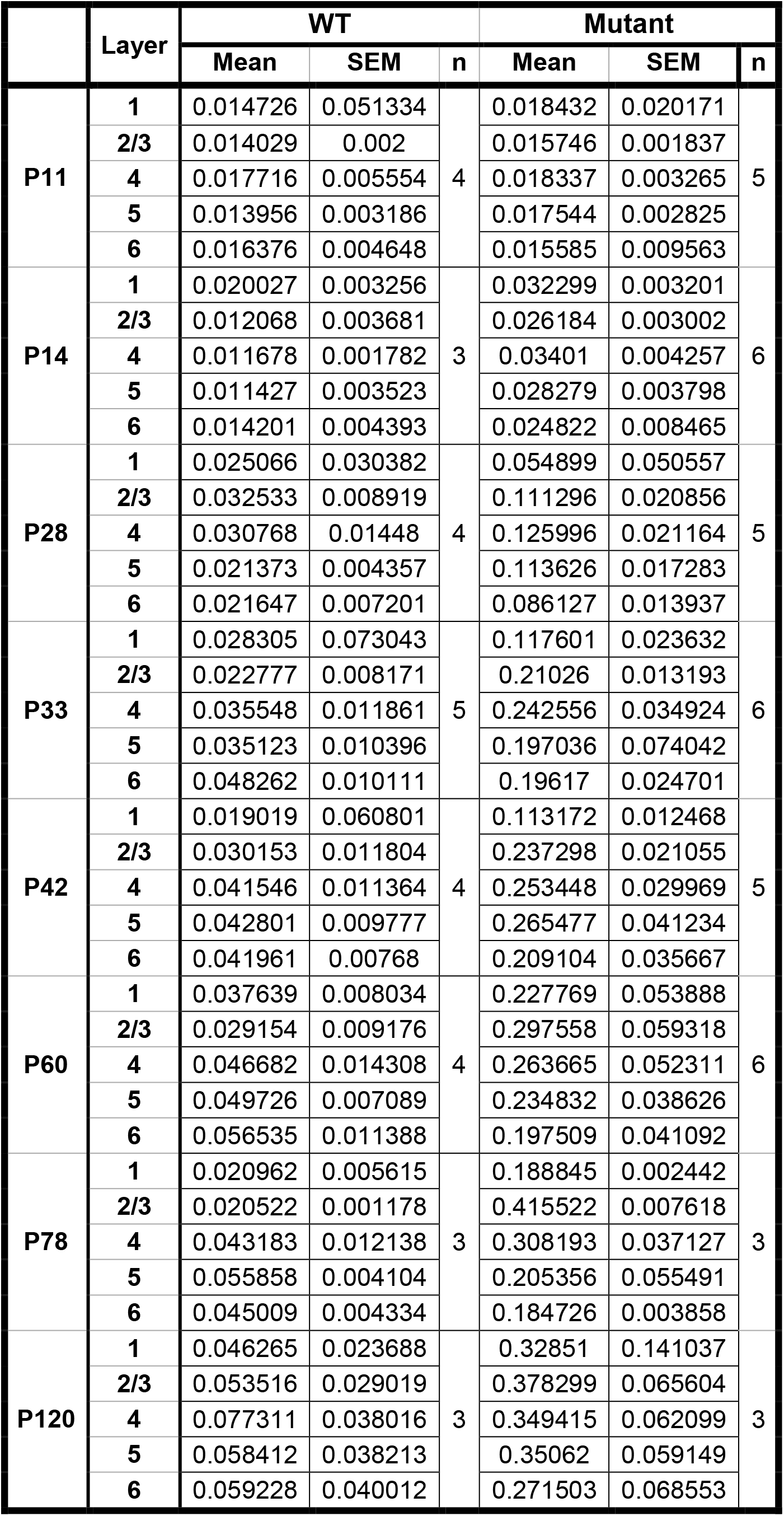
Mean values, s.e.m., and n for the bar chart in **Figure 1C**. Values are represented for each layer, at each age in WT and *Ppt1^-/-^* mice.

**Supplementary Figure 1.**
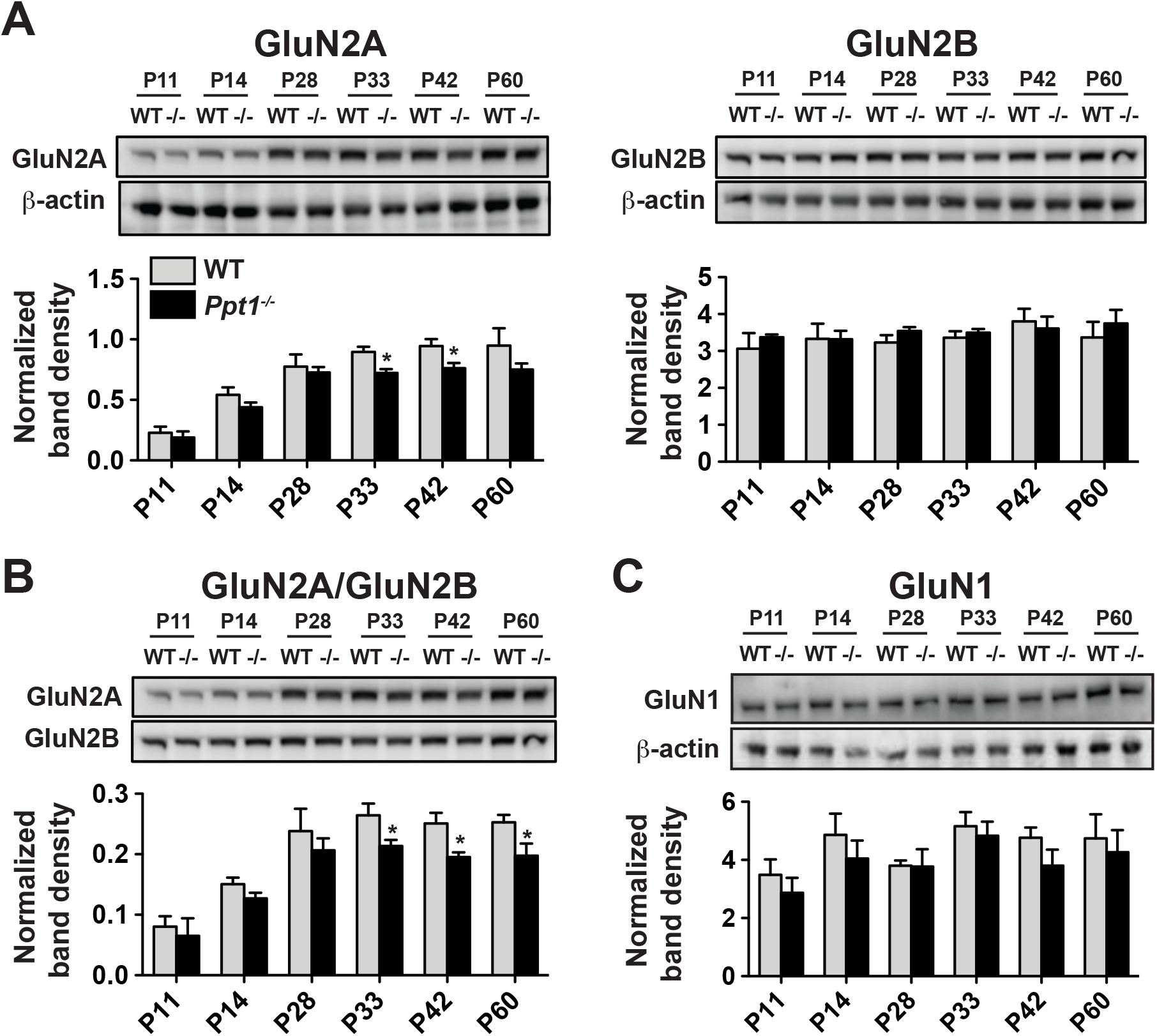
**(A)** Representative immunoblots of the GluN2 subunits, GluN2A and GluN2B in whole lysates across age and genotype as indicated (top) and quantification of band density (bottom) normalized to β-actin loading control within lane. **(B)** Representative immunoblots of GluN2A and GluN2B (top) from whole lysates across age and genotype and quantification of the ratio of GluN2A/GluN2B band density within animal (bottom). **(C)** Representative immunoblots of GluN1 in whole lysates across age and genotype as indicated (top) and quantification of band density (bottom) normalized to β-actin loading control within lane. For all experiments in **Supplementary Figure 1**, *Ppt1^-/-^* and WT were compared (n=4 independent experiments/animals with 2 repetitions/group) at each age using t-test and the significance was indicated as follows: *p<0.05. Error bars represent s.e.m.

